# Understanding range-wide immune gene variation in an endangered big cat (*Panthera tigris*)

**DOI:** 10.1101/2024.08.09.607326

**Authors:** B.V. Aditi, Uma Ramakrishnan

## Abstract

Assessing genetic diversity at loci putatively involved in fitness, such as immune genes, could provide important insights into the survival and vulnerability of endangered species. Research on immune genes in wild species has traditionally focussed on the Major Histocompatibility Complex (MHC) genes, which represent a small proportion of the immune gene repertoire and limits our understanding of immunity in non-model species. Here, we investigate five families of non-MHC immune genes - Tumor Necrosis Factors, interleukins, Toll-like Receptors, Leukocyte Immunoglobulin-Like Receptors, and chemokines - involved in adaptive and innate immunity in tiger *Panthera tigris*, an endangered carnivore known to have experienced historical bottlenecks and inbreeding. By leveraging 107 publicly available whole genomes, this study provides a comparative, multi-subspecific assessment of non-MHC immunogenetic diversity in tigers and identifies populations of conservation concern.

We found higher genetic variation at immune receptor genes compared to neutral regions and immune signaling genes. Site frequency spectra suggest balancing selection on Toll-like receptor genes. Further, tiger populations with histories of bottlenecks and inbreeding, such as South China tigers and North western Indian tigers exhibit reduced genetic diversity and higher mutation load at immune genes, compared to large and connected tiger populations. Overall, we find that genetic drift is likely the predominant evolutionary force shaping genetic variation in tigers at both neutral and immune loci. However, some immune polymorphisms persist despite drift, thereby maintaining elevated variation, indicating that tigers have not been entirely stripped of their immunogenetic resilience. Such immunogenetic resilience may prove vital as spillover threats intensify.

**Significance statement:** Understanding immune gene diversity beyond the MHC genes is essential for accurately assessing the adaptive potential of endangered species. Here, we leverage whole-genome sequencing across multiple tiger subspecies to show that non-MHC immune receptor genes harbour greater variation than neutral loci, and that balancing selection might maintain diversity at Toll-like receptor genes. Even in bottlenecked populations, some immune polymorphisms are retained, indicating that tigers possess broader immunogenetic resilience than previously thought.

## Introduction

Anthropogenic habitat loss confines wildlife into small and often isolated populations where genetic drift and inbreeding erode genetic variation and elevate mutation load (Frankham 1996; Keller and Waller 2002; Haddad et al. 2015; Grossen and Ramakrishnan 2024; Hasselgren et al. 2024). Hence, conservation geneticists have typically relied on genetic correlates of endangerment, such as genome-wide mutation load and genetic diversity, to inform and prioritise conservation efforts (Willi et al. 2022; Grossen and Ramakrishnan 2024). While these metrics are informative, there is a growing interest in characterising genetic diversity and mutation load at loci associated with fitness. Such variation has direct consequences for demographic outcomes such as survival and reproduction, and could provide complementary insights into population viability (Flanagan et al. 2018; Mable 2019). Since functional variation is shaped by distinct selective pressures in addition to genetic drift, its diversity dynamics are likely to differ from those of the rest of the genome.

Among fitness-related loci, immune genes are of particular interest as they sit at the interface of host-pathogen coevolution-a dynamic selective environment where genetic diversity could be depleted or maintained, depending on the mode of selection (Ejsmond and Radwan 2015; Ebert and Fields 2020). Like other loci, however, immune gene diversity can also be eroded through demographic processes such as bottlenecks and drift, and such loss is associated with increased disease vulnerability (Martin et al. 2022; Cortazar-Chinarro et al. 2022). Landmark studies on cheetahs and Tasmanian devils have established links between genetic erosion, immune gene variability and pathogen susceptibility. Several studies find that immune genetic diversity in bottlenecked populations reflects that at neutral loci (Bollmer et al. 2011; Strand et al. 2012a; Mathur et al. 2023) but other studies observe higher polymorphisms in genes under balancing selection, preventing a complete erosion of genetic diversity at these loci even in such vulnerable populations (Edwards et al. 2000; Gilroy et al. 2017; Schubert et al. 2025). This raises the question of whether selection can buffer immune loci against demographic erosion. Consequently, understanding genetic variation at immune-related genes in wild populations could provide crucial information for understanding disease susceptibility and resistance.

Most studies on population genetics of immune genes focus on a single family-the Major Histocompatibility Complex (MHC), which is involved in adaptive immunity (for e.g. Aguilar et al. 2004; Castro-Prieto et al. 2010; Silver et al. 2024) but MHC genes represent a small proportion of the immune gene repertoire – approximately 1% of ∼6,000 genes of the immunome (Acevedo-Whitehouse and Cunningham 2006; Bhattacharya et al. 2018). Hence, attention is expanding towards other immune gene families to obtain a more holistic view of immunogenetic variation. Genes for Toll-Like Receptor (TLR) and chemokines are characterized in several wild species, including those that exist in small populations (for e.g. Turner et al. 2011; Morris et al. 2015; Vlček et al. 2023). Genes that code for receptors directly interacting with pathogen molecules (e.g., MHC and TLR genes) show signatures of balancing or diversifying selection. In contrast, genes that code for signalling molecules, such as interleukins and chemokines show signatures of purifying selection (Mukherjee et al. 2009). Moreover, immune genes are often examined within a single population or species, making it challenging to establish a reliable baseline for assessing genetic metrics (Bosse and van Loon 2022). We address these limitations by studying five non-MHC immune genes in a comparative framework across six tiger subspecies.

Tigers (*Panthera tigris*) are a flagship endangered species that have undergone population and range decline due to large-scale poaching and habitat loss, and have low genome-wide genetic diversity (Mondol et al. 2013; Armstrong et al. 2021). Some populations are recovering demographically (Jhala et al. 2025), but they face several threats, including exposure to infectious diseases (Sidhu et al. 2019) which could be compounded by increasing density in small populations (Gilbert et al. 2023a). Tigers have been mainly known to be affected by viral infections such as severe acute respiratory syndrome coronavirus 2 (SARS-CoV 2) and Canine distemper virus (CDV), followed by bacteria, protozoa and helminths (McCauley et al. 2021; see review in Gilbert et al. 2023a). Non-MHC genes such as TLR, TNF and other components of the innate immune system are known to be involved in immunity against these pathogens (Areal et al. 2011; Capozza et al. 2021; de Almeida Lopes et al. 2025). Although our knowledge about direct links between disease and immune genes in tigers remains poor, studies on other felids indicate inbreeding and loss of genetic diversity at immune genes could have effects on disease susceptibility (e.g. cheetahs and Tasmanian Devil). Notably, inbred Amur tigers exhibit reduced immune heterogeneity and dysregulated immune signaling pathways associated with inflammation and cancer, suggesting that inbreeding has tangible phenotypic consequences for immune function (Bi et al. 2026). Investigating the immunogenetic variation of range-wide tigers could improve our understanding of fitness-related genetic diversity and the relative vulnerability or resilience of endangered populations.

Globally, there are six extant tiger subspecies/populations recognized based on genetic and phenotypic data: Bengal tiger *Panthera tigris tigris* (Linnaeus, 1758), Sumatran tiger *P.t.sumatrae* (Pocock, 1929), Indochinese tiger *P.t. corbetti* (Mazak, 1968), Malayan tiger *P.t.jacksoni*, Amur tiger *P.t. altaica* (Temminck, 1844) are subspecies existing in the wild, while South China tiger *P.t. amoyensis* (Hilzheimer, 1905) is extinct in wild but survive in captivity (Figure 1). Within India, tigers exhibit strong genetic differentiation, resulting in at least three subpopulations: the North West (NW) Indian subpopulation is small and isolated, while the Central Indian (CI) and South Indian (SI) subpopulations are large and interconnected (Natesh et al. 2017; Armstrong et al. 2021) (Figure 1). The Northwest Indian subpopulation shows genomic signatures of inbreeding (Khan et al. 2021). Studies on MHC genes in tigers have revealed signatures of positive, balancing, and purifying selection, and also reveal low allelic diversity compared to humans or domestic cats (Pokorny et al. 2010; Wei et al. 2010; Shukla et al. 2023). Currently available whole-genome resequencing data for various tiger subspecies offer a unique opportunity to investigate immunogenetic variation at a broader scale, including multiple subspecies and their subpopulations, particularly those with known histories of disease exposure such as Amur tigers that experience Canine Distemper Virus epidemics (Gilbert et al. 2023c) and Northwest Indian tigers who are exposed to CDV through domestic dogs in their surroundings (Sidhu et al. 2019).

**Figure 1:**
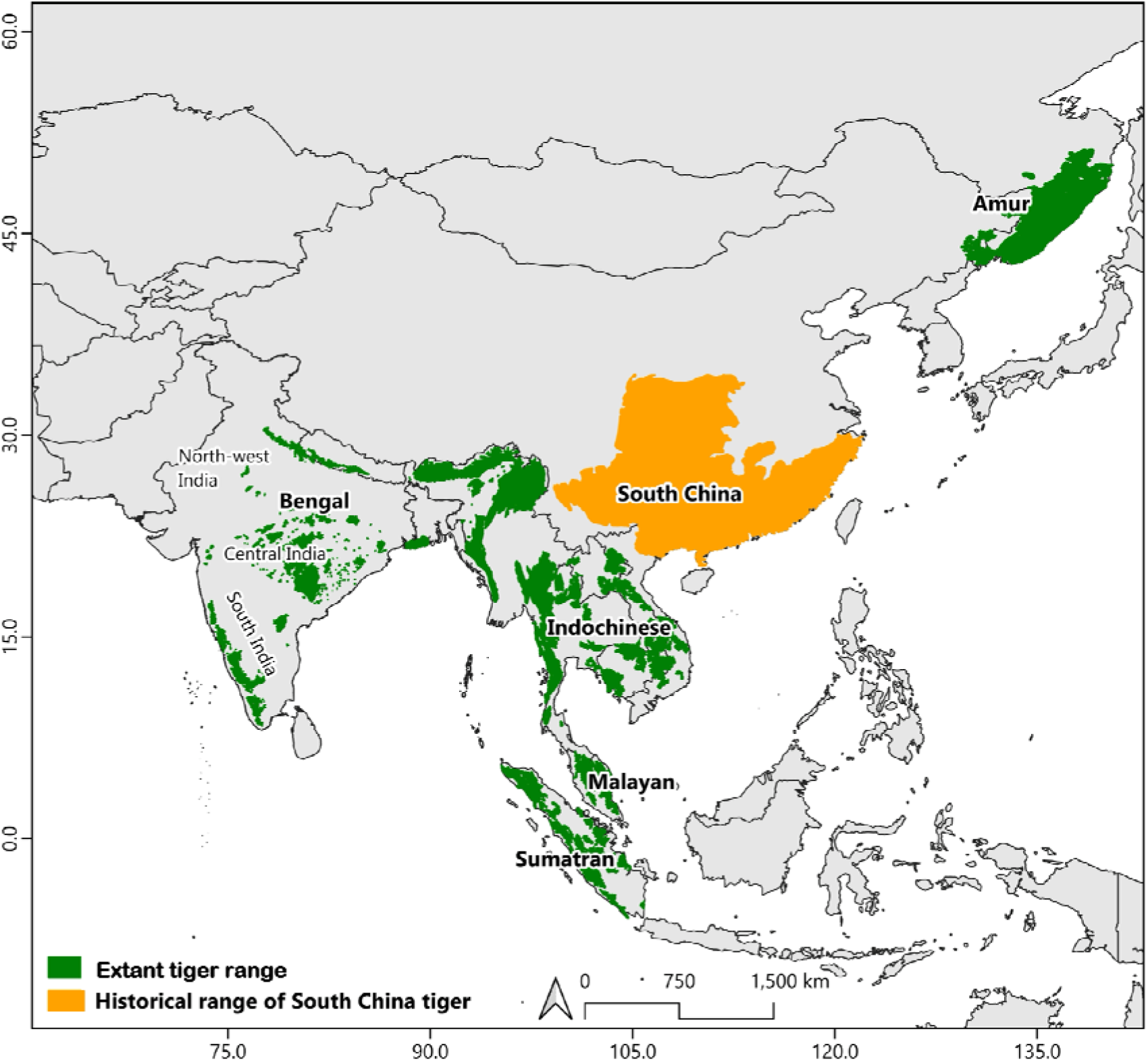
Extant and historical distribution of tiger (*Panthera tigris*) subspecies and Indian subpopulations analyzed in this study.

In this paper, we examine: i) Genetic variation and differentation at immune and neutral loci across a) five non-MHC gene families and b) different tiger populations (six subspecies as well as three Indian sub-populations), ii) Site frequency spectra of immune gene loci for signatures of selective pressures, iii) Putatively deleterious variation at immune genes. The immune genes we analyzed were examined individually but were categorized based on their functional roles to facilitate biologically meaningful interpretation and increase inferential robustness. This framework offers a conservative approach for inferring selection as genes within a functional class are more likely to experience similar selective pressures. Genes encoding pathogen recognition molecules, typically receptor proteins, were considered separately from genes encoding signaling molecules such as cytokines. This framework allows us to establish species-specific baselines for each immune gene category, enabling comparisons of diversity patterns within species. Finally, using these metrics, we identified populations of conservation concern based on accumulated mutation load and lower genetic diversity.

## Results

### Summary of Immune genes identified

We were able to identify 91 Immune Genes spanning 11,57,251 bp across five gene families related to immunity with complete and contiguous gene sequences. Two gene families code for receptor molecules that are involved in adaptive and innate immunity. Toll-like receptor (TLR) genes code for receptor molecules that recognize pathogens as part of the innate immune mechanism. Leukocyte-like Immunoglobulin-Like Receptor (LILR) molecules interact with MHC receptors and regulate the activity of NK Cells and are part of the adaptive immune response. The TNF, Interleukin, and chemokine gene families code for small, secreted molecules called cytokines and their ligands involved in innate immunity, which reach sites of immune reactions and aid in the process by inducing inflammation and signaling. Table 1 presents the number of genes, their sizes, and the number of SNPs across the entire gene in these gene families. The Supplementary Table 2 lists the names of the genes within each family.

**Table 1:** Compilation of immune gene families identified from range-wide tiger genomes.

| Gene Family | Number of Genes | Total length in kb | Number of SNPs | Function |
| --- | --- | --- | --- | --- |
| **Genes coding for receptors** |  |  |  |  |
| Leucocyte Immunoglobulin Like Receptor (LILR) | 6 | 85,394 | 269 | Expressed on macrophages and dendritic cells, and regulates the killing activity of the cells interacting with MHC Class I molecules |
| Toll-Like Receptor (TLR) | 5 | 50,969 | 114 | Receptors on Macrophages and dendritic cells, that recognize structurally conserved molecules derived from microbes |
| **Genes coding for signalling molecules** |  |  |  |  |
| Tumor Necrosis Factor (TNF) | 32 | 4,49,348 | 1076 | A cell signalling protein (cytokine) involved in systemic inflammation, and is one of the cytokines that make up the acute phase reaction. |
| Chemokine | 41 | 3,10,833 | 687 | Receptors on the surface of leucocytes and cytokines are involved in the immune response |
| Interleukin | 7 | 2,60,707 | 584 | Secreted proteins involved in the regulation of the immune system |

### Patterns of genetic diversity at immune genes in tigers

To assess patterns of genetic diversity at immune genes and neutral loci, we estimated nucleotide diversity over 500 bp windows and heterozygosity for each SNP. Receptor loci showed elevated nucleotide diversity relative to neutral and signalling loci, while signalling loci showed the lowest diversity of the three categories. Mean nucleotide diversity at receptor loci (mean=0.0016, 95 % CI: 0.00047-0.0033, GLM estimate = -566.21, p-value=<2e-16) is significantly higher than neutral loci (mean=0.0007, 95 % CI: 0.0007-0.0007, GLM estimate= 806.77) (Figure 2a), whereas signalling loci show the lowest mean nucleotide diversity (0.0003, 95 % CI: 0.0003-0.0004, GLM estimate= -3.82, p-value=0.91). Heterozygosity patterns did not differ significantly between locus categories. Mean heterozygosity at receptor loci (0.174, 95 % CI: 0.143- 0.206, pseudo-R^2^=7.97e-06,p-value=0.43) and signalling loci (0.149, 95 % CI: 0.136-0.163, pseudo-R^2^ =7.97-06, p-value = 0.0674) is lower than neutral loci (0.2, 95 % CI: 0.2 - 0.2) (Figure 2b) but this difference is not significant.

**Figure 2:**
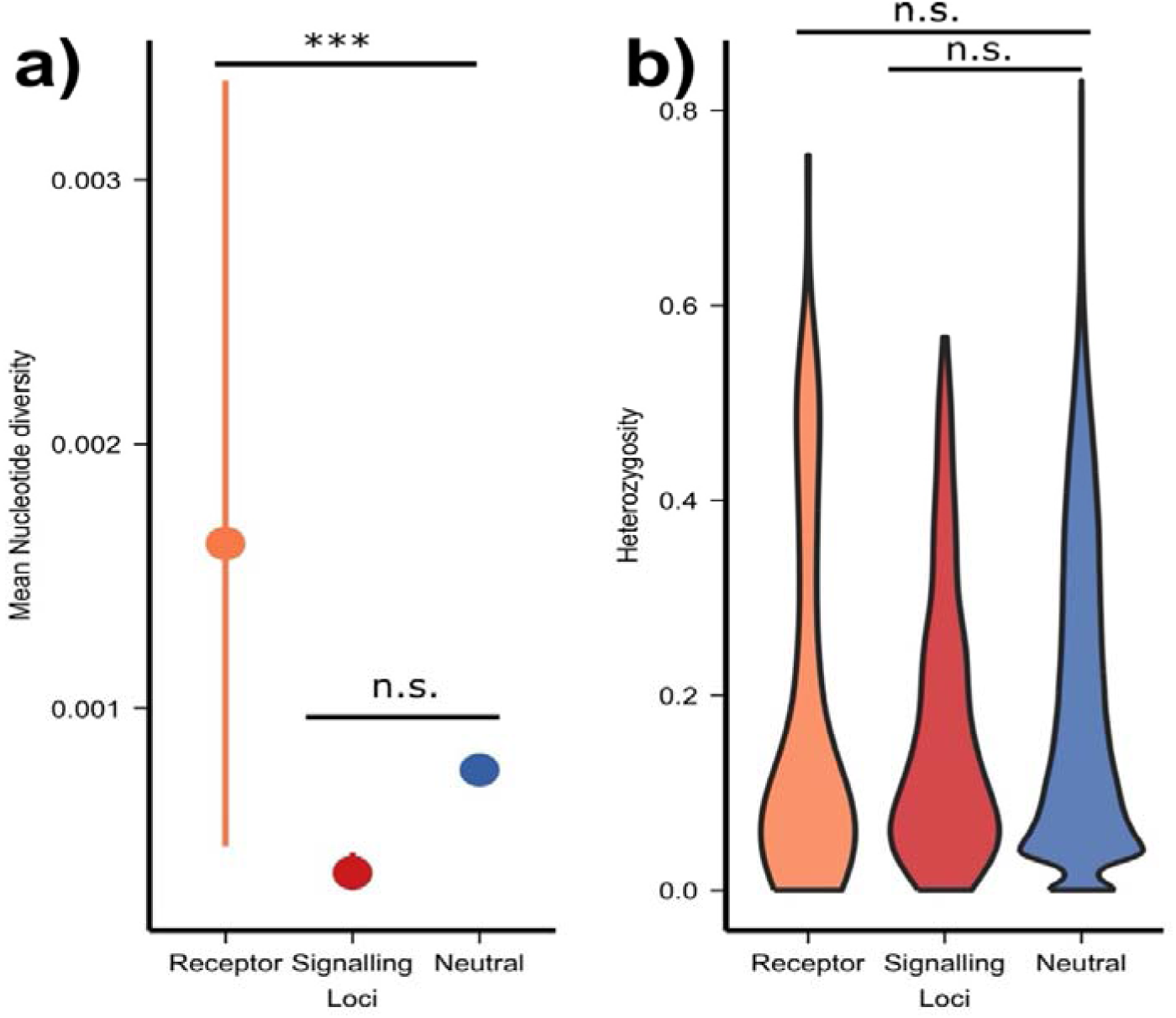
Distribution of a) nucleotide diversity (with 95% confidence intervals around the mean and b) heterozygosity at receptor and signalling immune genes and neutral loci (Significance levels: *** < 0.000, ** < 0.001, * < 0.01, ns < 0.05) **Alt text:** Graph a) shows mean and confidence intervals of immune receptor and signalling genes and b) shows violin plots of heterozygosity

We also examined the distribution of nucleotide diversity and heterozygosity values at each family of immune genes (Supplementary Table 3). We found significant differences in nucleotide diversity of TLR (mean=0.0021, 95 % CI: 0.0004-0.005, GLM estimate = -584.58, p-value=<2e-16) and LILR genes (mean=0.001, 95 % CI: 0.0002-0.0028, GLM estimate = -542.55, p-value=<2e-16) compared to neutral loci, but no significant differences in heterozygosity were observed (Figure 3, Supplementary Tables 4 and 5). Nonetheless, we observe a higher mean nucleotide diversity and higher (non-significant) heterozygosity at the receptor genes (LILR and TLR) (Figure 3a, Figure 3b).

**Figure 3:**
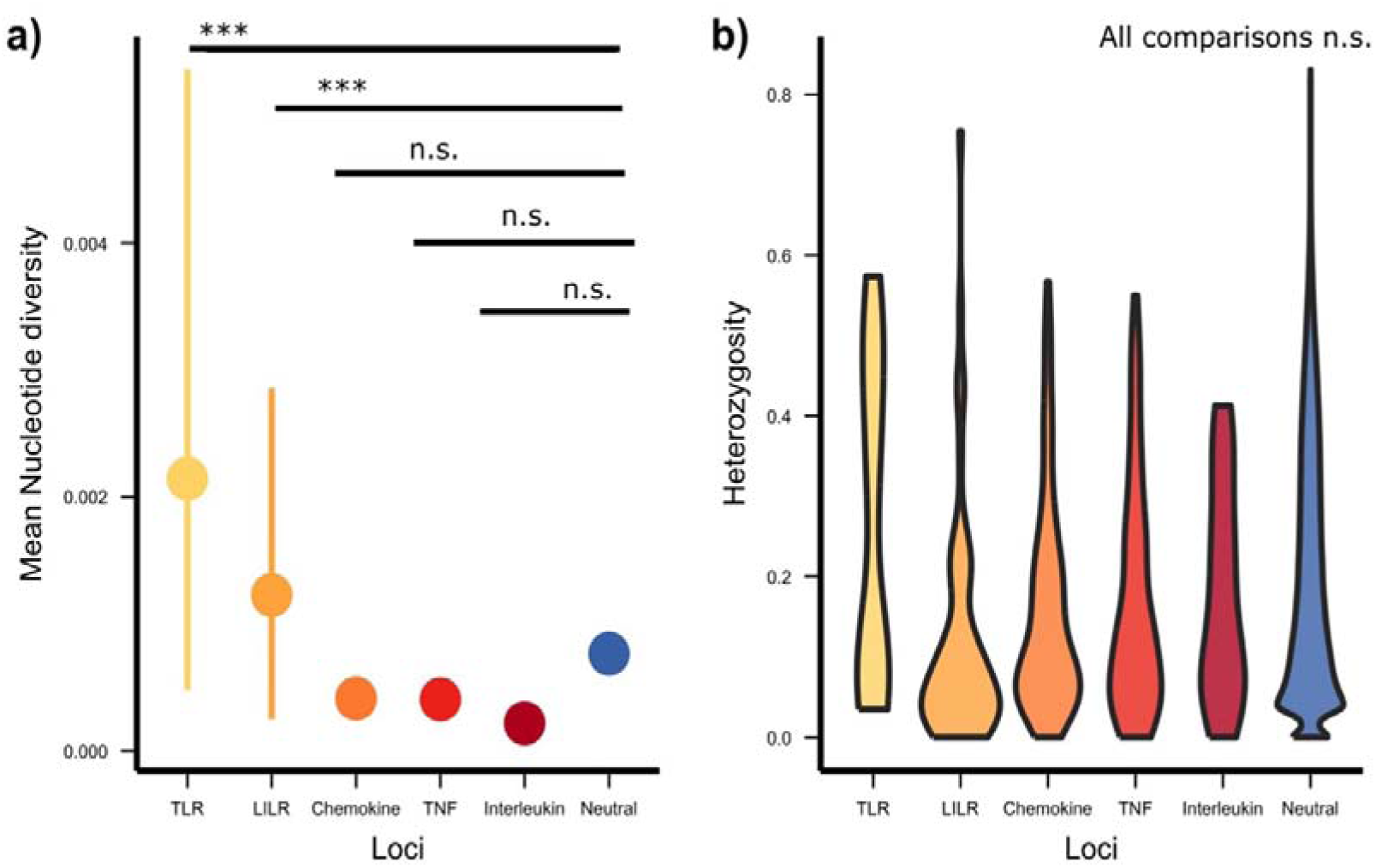
Distribution of a) nucleotide diversity (with 95% confidence intervals) (Significant difference from neutral in: TLR ***, LILR ***, chemokine, n.s, TNF n.s, interleukin n.s) and b) heterozygosity at different families of immune genes (No significant differences between groups) (Significance levels: *** < 0.000, ** < 0.001, * < 0.01, ns < 0.05) **Alt text:** Graph a) shows mean and confidence intervals of immune receptor and signalling genes and b) shows violin plots of heterozygosity

### Differences in immune gene diversity across subspecies and Indian subpopulations of tigers

We investigated the genetic diversity patterns for neutral and immune loci across range-wide subspecies. Since we found no significant differences in heterozygosity of different gene families, we combine them to contrast immune genes with neutral loci in this analysis. Nucleotide diversity at receptor loci differs significantly from that at neutral loci; therefore, we compare receptor and signalling loci separately with neutral loci.

Our results show that both nucleotide diversity (Figure 4, Supplementary Table 6) and heterozygosity (Figure 5, Supplementary Table 7) vary similarly across subspecies at both immune and neutral loci. While these differences were significant in most comparisons of nucleotide diversity, none of the differences were significant in comparisons of heterozygosity (detailed zero-inflated and binomial GLM results Supplementary Tables 8, 9, 10, and 11). Sumatran (SUM) tigers have the least nucleotide diversity at immune and neutral loci (Figure 4a), and South China (SOC) tigers have the least heterozygosity at immune and neutral loci (Figure 5a). Malayan (MAL) tigers exhibit the highest mean nucleotide diversity and heterozygosity at both immune and neutral loci (Figures 4a and 5a). In populations within India, the nucleotide diversity and heterozygosity at immune loci follow a pattern similar to neutral loci (Figures 4b and 5b). The large, connected populations (CI and SI) have higher nucleotide diversity and heterozygosity when compared to the small, isolated population (NW). The differences in distributions across populations, as tested by Kruskal-Wallis’s test, were found to be significant, except for nucleotide diversity of receptor loci in Indian subpopulations (Supplementary Table 12 and Supplementary Table 13).

**Figure 4:**
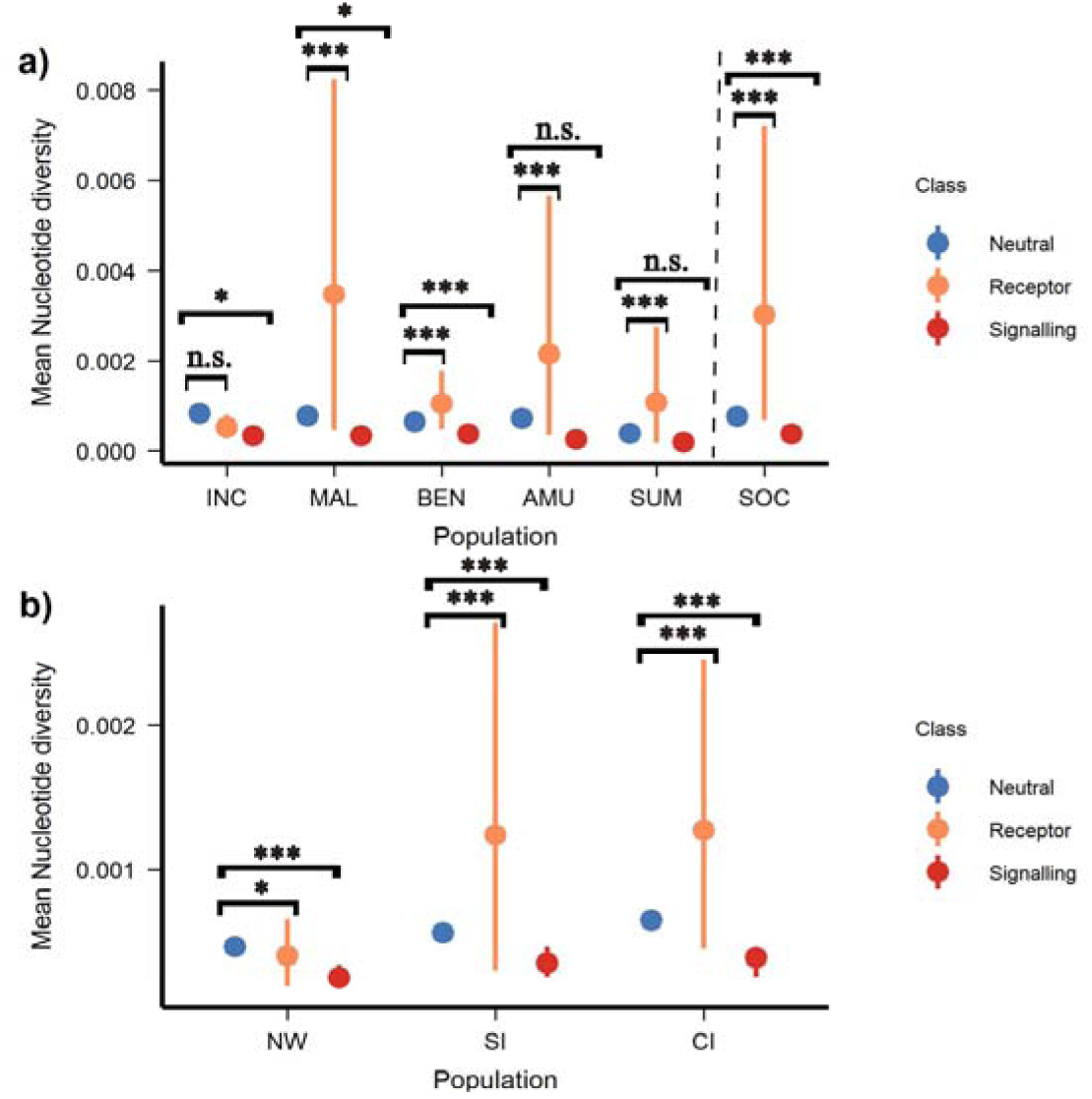
a) Mean and confidence intervals for nucleotide diversity at immune receptor, immune signalling, and neutral loci in extant subspecies of tigers, including INC (Indochinese), MAL (Malayan), BEN (Bengal), AMU (Amur), SUM (Sumatran) and SOC (South China). Dotted line separates the captive SOC population from wild populations. b) Mean and confidence intervals for nucleotide diversity at immune receptor, immune signalling, and neutral loci in Indian tiger populations NW (Northwest), CI (Central India), and SI (South India). (Significance levels: *** < 0.000, ** < 0.001, * < 0.01, ns < 0.05) **Alt text:** Graph a) shows mean and confidence intervals of nucleotide diversity of immune receptor and signalling genes in different tiger subspecies and b) shows mean and confidence intervals of nucleotide diversity of immune receptor and signalling genes in different Indian tiger subpopulations

**Figure 5:**
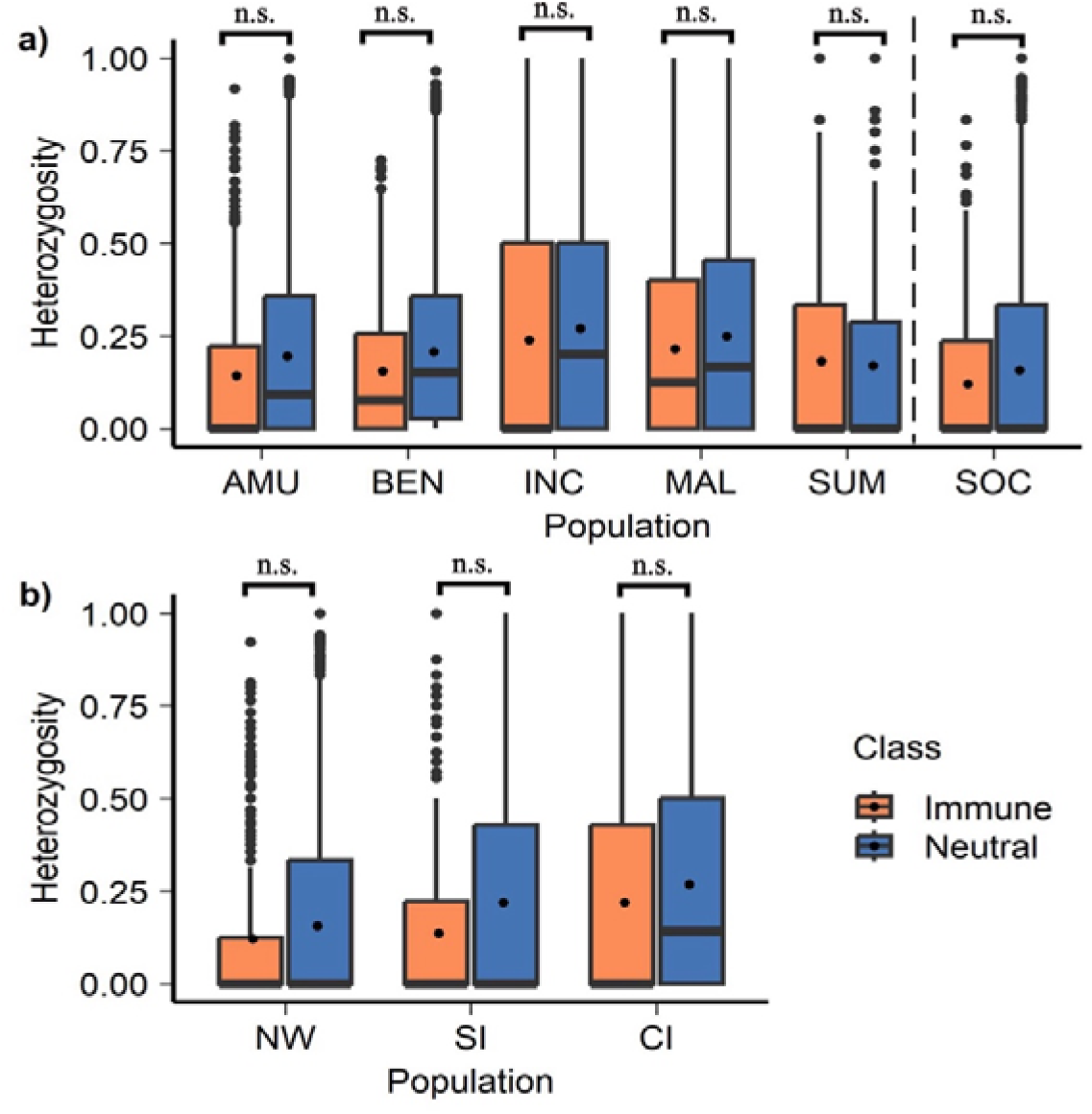
a) Box and whisker plots for heterozygosity at Immune and Neutral loci for extant subspecies of tigers, including INC (Indochinese), MAL (Malayan), BEN (Bengal), AMU (Amur), SUM (Sumatran) and, SOC (South China). Dotted line separates the captive SOC population from wild populations. (b) Box and whisker plots for heterozygosity at Immune and Neutral loci for different populations of Indian tigers NW(Northwest), CI (Central India) and SI (South India) (Significance levels: *** < 0.000, ** < 0.001, * < 0.01, ns < 0.05) **Alt text:** Graph a) shows boxplots of heterozygosity immune receptor and signalling genes in different tiger subspecies and b) shows boxplots heterozygosity of immune receptor and signalling genes in different Indian tiger subpopulations

Receptor loci exhibit consistently high nucleotide diversity for subspecies, as well as Indian subpopulations. The only exception is the Indochinese (INC) tiger, for which receptor immune nucleotide does not differ significantly from neutral nucleotide diversity values (Figure 4a). Signalling loci showed lower mean nucleotide diversity compared to neutral diversity in all populations examined; however, this difference was not significant in all comparisons as evident in Figure 4 (Supplementary Tables 8 and 9).

Although the mean and median values for both diversity indices were significantly lower in the Northwest Indian (inbred) population, several outliers were present in this distribution. We also examined the outliers in the other populations with low genetic diversity i.e. in the Amur, Sumatran and South China tiger populations. We found that the outlying SNPs were from sites across receptor and signalling genes and were not restricted to any one gene family and outliers in different populations were in the same few genes (listed in Supplementary Table 14).

### Genetic differentiation at immune gene and neutral loci between populations of tigers

Given the differences in habitats (and associated pathogens) of the different subspecies and subpopulations of tigers, differential pathogen pressure may lead to genetic differentiation at immune loci. We compared genetic differentiation at immune loci to that at neutral loci using Hudson’s F_ST_ for tiger subspecies (Figure 6a) and Indian subpopulations (Figure 6b). We found no significant difference in F_ST_ distributions between immune receptor (p-value= 3.46e-11), signalling (p-value= 0.33), and neutral loci (Gaussian GLM, pseudo-R^2^ = 2.74e-06) at subspecies level as well as at subpopulation level (receptor p-value= 0.7948, signalling p-value= 0.0098 and Gaussian GLM, pseudo-R^2^ = 0).

**Figure 6:**
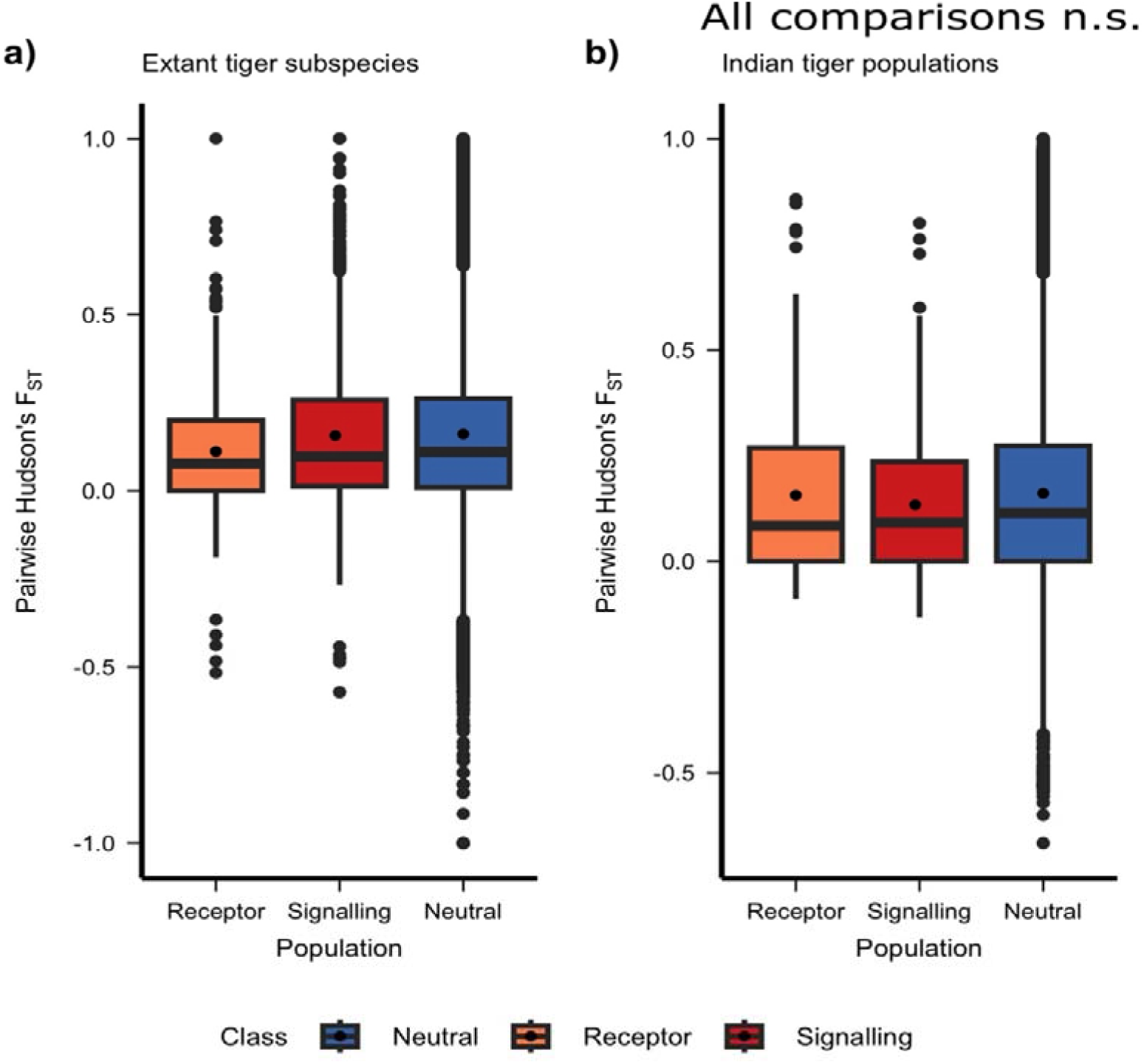
a) Box and whisker plots for the fixation index at the immune receptor, immune signalling, and Neutral loci for different subspecies of tigers. b) Box and whisker plots for the Fixation index at immune receptor, immune signalling, and neutral loci for different populations of Indian tigers (No significant differences found) (Significance levels: *** < 0.000, ** < 0.001, * < 0.01, ns < 0.05) **Alt text:** Graph a) shows boxplots of immune receptor and signalling genes in different tiger subspecies and b) shows boxplots of immune receptor and signalling genes in different Indian tiger subpopulations

We also examined F_ST_ at immune and neutral loci for populations that have a history of disease or drastic demographic change as both scenarios could lead to elevated F_ST_ values. We compared F_ST_ for Amur, Sumatran, and South China tiger populations with all other tiger populations and the three Indian subpopulations, and found no significant differences. Sumatran tigers had the highest mean F_ST_ at both neutral and immune loci. Amur tigers had slightly higher mean F_ST_ at immune loci when compared to neutral loci. Northwestern Indian tigers were the most genetically differentiated amongst the three Indian tiger populations, and similar patterns were reflected in immune genes with no significant difference between neutral and immune loci (Supplementary Figure 1 and Supplementary Figure 2, Gaussian GLM Supplementary Table 15).

### Comparison of site-frequency spectra between immune and neutral loci

Unfolded SFS for synonymous and nonsynonymous variants were found not to differ significantly from neutral SFS (Supplementary Table 16). However, the proportion of nonsynonymous SNPs at lower frequencies was higher than that of neutral or synonymous SNPs, which could indicate weak purifying selection (Figure 7). Since TLR and LILR genes have significantly higher nucleotide diversity than neutral loci, we assessed selective pressure at these loci through SFS of synonymous and non-synonymous SNPs (Figure 8). We found that the SFS of TLR genes differed from that of neutral SFS, with more non-synonymous variants at intermediate frequencies; however, this difference was not statistically significant (Supplementary Table 16). The SFS of LILR genes was not different from neutral SFS. The TLR genes identified to harbor SNPs with high heterozygosity were TLR2, TLR5, and TLR9.

**Figure 7:**
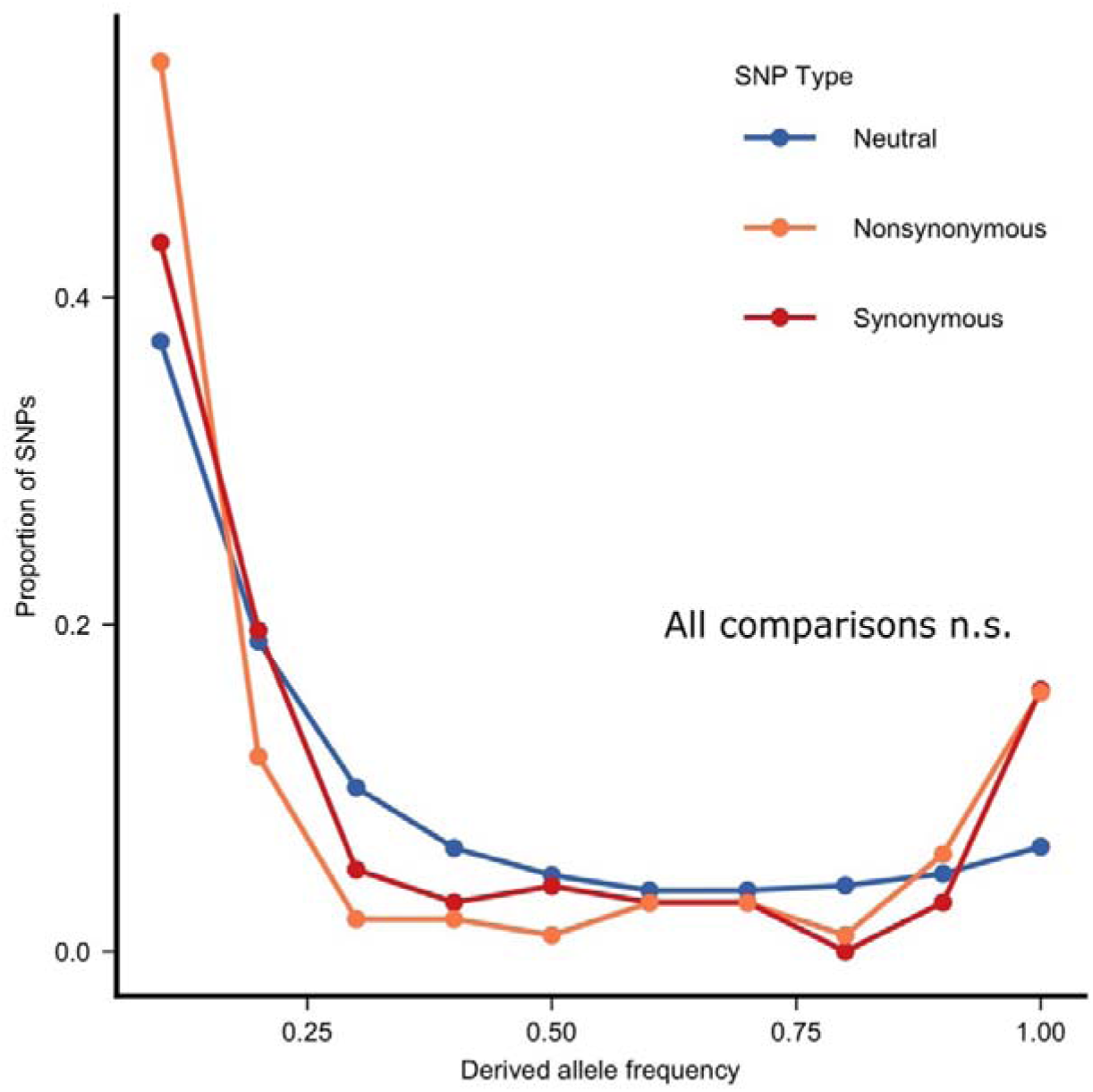
Comparison of SFS of neutral SNPs with that of immune synonymous and nonsynonymous SNPs (Significance levels: *** < 0.000, ** < 0.001, * < 0.01, ns < 0.05) **Alt text:** Graph of site frequency spectra of immune synonymous, nonsynonymous and neutral SNPs

**Figure 8:**
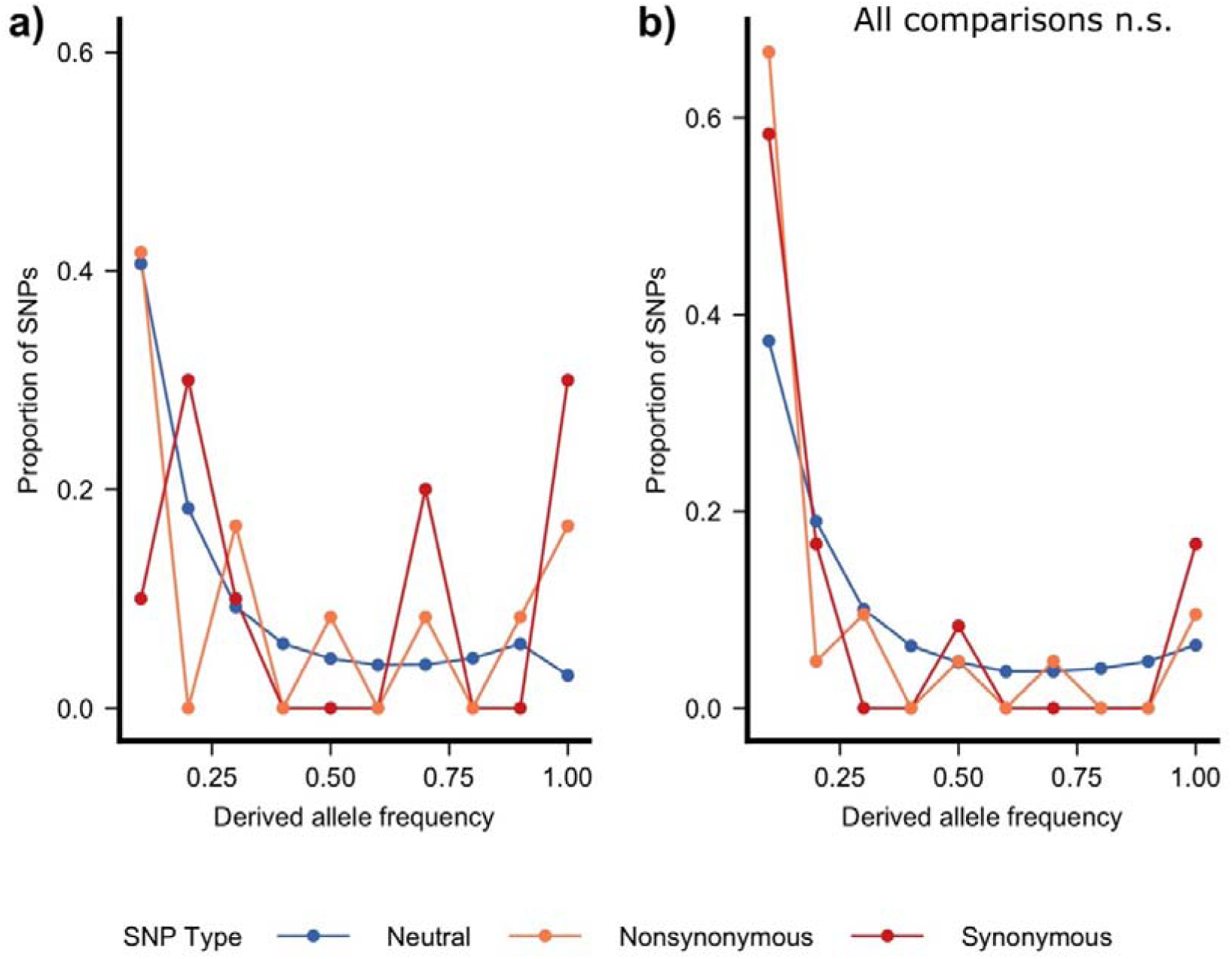
Comparison of SFS of neutral SNPs with that of synonymous and nonsynonymous SNPs at a) TLR genes and b) LILR genes (no significant differences found) (Significance levels: *** < 0.000, ** < 0.001, * < 0.01, ns < 0.05) **Alt text:** Graph of site frequency spectra of immune synonymous, nonsynonymous and neutral SNPs at a) TLR and b) LILR

### pDeleterious and pLOF mutation load in Indian populations

The pDeleterious and pLOF mutations identified from VEP and further confirmed through Polyphen (Supplementary Figure 3) were used to estimate mutation load in the Indian subpopulations. The majority of the mutation load was found to be masked (Figure 9). We found that large, connected populations examined (SI, CI, and MAL) have a higher masked pDeleterious load than the inbred populations (NW and SOC). Inbreeding load, estimated as homozygous damaging alleles, was lower than masked load in all populations for both pDeleterious and pLOF. The two inbred populations harbored pLOF load (both masked and inbreeding), but SOC tigers lacked any homozygous deleterious mutations (Figure 9).

**Figure 9:**
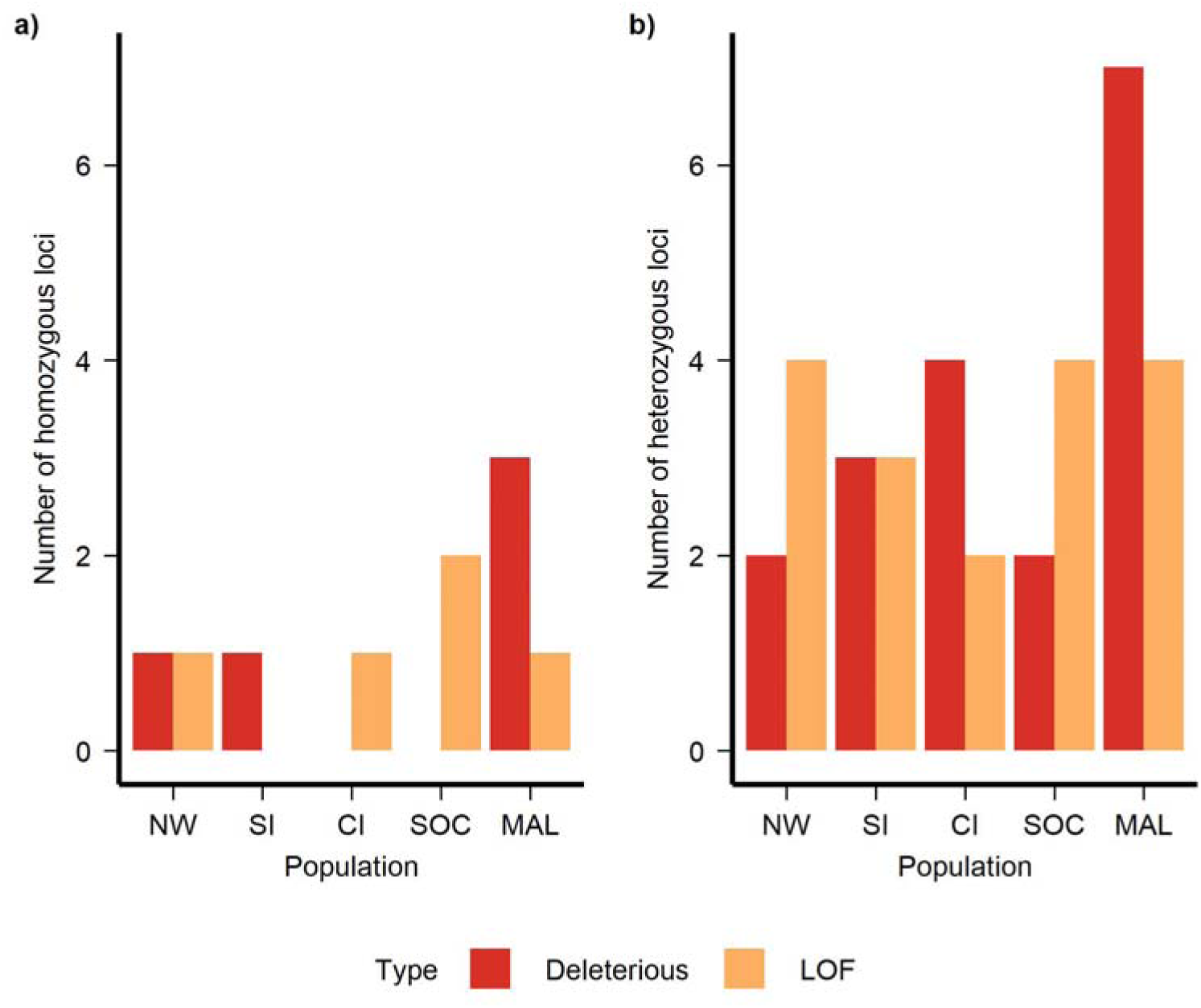
a) Homozygous and b) masked burden of pDeleterious and pLOF (Loss of Function) load in small, isolated inbred NW (Northwest) population and large, connected populations SI (South India) and CI (Central India); inbred South China (SOC) and population with high genetic diversity MAL (Malayan) tigers **Alt text:** Barplots showing mutation load of pDeleterious and pLOF in different tiger populations Homozygous pDeleterious mutations were present in CCR2 (MAL), CCL8 (MAL), and TLR5 (NW and SI). The homozygous pLoF mutations were identified in CCL5 (CI, MAL, and SOC), CXCR2 (SOC), and TNFSF11 (NW).

## Discussion

### Heterozygosity and nucleotide diversity at different immune gene families reveal contrasting patterns

We characterized genetic variation at several families of immune genes in tigers spanning adaptive and innate immune functions. Given the availability of gene annotations and genomic data, we examined a total of 91 genes spanning 11,57,251 bp, as opposed to a few genes and shorter stretches studied through molecular methods such as PCR (Pokorny et al. 2010; Morris et al. 2015). Using genetic diversity indices of nucleotide diversity and heterozygosity at intergenic regions, we inferred neutral diversity, against which we compare the same diversity indices for immune genes. Immune genes that code for receptors exhibited significantly higher nucleotide diversity than signalling immune and neutral loci. TLR genes also exhibit higher heterozygosity, although this difference is not statistically significant. However, immunogenetic variation largely falls within neutral expectations suggesting the major role of genetic drift. The receptor and signaling genes differ in total length and number of SNPs, but our analytical approach accounts for this in several ways. Heterozygosity was calculated per SNP and nucleotide diversity was estimated within fixed 500 bp windows, normalising for differences in gene length. The consistently elevated diversity in the receptor gene across both metrics is therefore unlikely to reflect a methodological artifact of gene size differences, and is more consistent with biological variation.

### Patterns of genetic diversity of immune genes mirror that of neutral loci across populations with differing demographic histories

To further assess the effect of drift as a dominant evolutionary force, we compared immune and neutral variation across tiger subspecies and among Indian tiger populations that differ in in terms of demographic history. We find that patterns of immune variation across populations mirror neutral variation, confirming that most immune genes examined are shaped largely by genetic drift. While immune receptor genes are consistently more diverse than neutral loci within populations, diversity patterns across populations parallel neutral diversity. Our results substantiate inferences from earlier studies that most loci in the coding regions of the tiger genome are influenced by drift, regardless of the subspecies or population, potentially due to demographic fluctuations and bottlenecks (Armstrong et al., 2021).

The Malayan, Indochinese, and Bengal tigers (Figures 3a and 4a) were found to have the highest genetic diversity at both immune and neutral loci, consistent with their proximity to the ancestral population and the large size of the Bengal population. Sumatran tigers, being an island population, have the lowest nucleotide diversity at immune and neutral loci (Clegg et al. 2002). The South China tigers, which are extinct in the wild and exist as a captive population, were found to have the lowest heterozygosity at immune and neutral loci, as expected in an inbred population. Amur tigers, having undergone strong recent bottlenecks similar to those of Sumatran tigers, exhibit genetic diversity comparable to that of Sumatran tigers at both immune and neutral loci.

Our results corroborate findings in several other endangered species, where a decline in population size and isolation leads to loss of adaptive genetic diversity, along with neutral diversity. Such patterns have been found for immune gene diversity in Tasmanian devils (Morris et al. 2015), black grouse (Strand et al. 2012b), dunnocks (Lara et al. 2020), snub-nosed monkeys (Luo et al. 2012).

### Several immune genes retain high polymorphisms even in bottlenecked populations

We observe several genes with high heterozygosity values in the Northwestern population of Indian tigers, which is a small, isolated, and inbred population. This is also true in Sumatran, South China, and Amur tigers (Supplementary Table 13). Importantly, these genes represent a broad spectrum of immune functions, rather than being restricted to receptor-mediated ones. Overall, this suggests that despite a bottleneck and drift, some loci might retain high polymorphisms, potentially maintained by past selection. Such variation has been described previously as “ghosts of selection past” and may be eroded over time due to drift debt (Oliver and Piertney 2012; Gilroy et al. 2017; Davies et al. 2021). Interestingly, a previous study on MHC Class I and Class II genes in Asiatic lions and Indian tigers found ‘hyper polymorphic’ hotspots even when allelic diversity was low (Sachdev, 2005; Pokorny et al. 2010).

### Patterns of genetic differentiation at immune loci in tigers are similar to differentiation at neutral loci

We used the fixation index, or F_ST_, to contrast differentiation at the subspecies level and among Indian populations with differentiation at neutral loci. F_ST_ patterns reiterate the predominant role of drift in shaping the evolution of tigers, including immune loci. F_ST_ values for population pairs, where one (a) had a history of disease (e.g. Amur tigers (Gilbert et al. 2014)), (b) had undergone major bottlenecks (e.g. South China and North western Indian tigers), or (c) had undergone founding events (e.g. Sumatran tigers in the Sunda islands) were not significantly different for immune gene and neutral loci. The largely solitary ecology of tigers, with limited social contact among conspecifics and other felids, likely restricts horizontal pathogen transmission and weakens selective pressures on immune genes. Oral consumption and nasal exposure could be two possible modes of infection. This could have prevented a high prevalence of infections and weaker pathogen-mediated selection, which would be difficult to detect over the strong signatures of demographic history.

### TLR genes show signatures of balancing selection

The high genetic diversity in TLR genes could be a result of balancing selection. We compared SFS for neutral and immune synonymous and nonsynonymous SNPs to detect signatures of balancing selection while accounting for a background of demographic history (Weedall and Conway 2010; Fijarczyk and Babik 2015; Soni and Jensen 2024). We found higher frequencies of nonsynonymous SNPs at intermediate frequencies in TLR genes but not in LILR genes. Such a signal is associated with balancing selection in previous studies (Bernhardsson and Ingvarsson 2011; Fijarczyk et al. 2016; Bitarello et al. 2018; Brandt et al. 2018). After MHC loci, TLR loci are frequently found to be under balancing selection (Kloch et al. 2018). TLR genes are involved in the recognition of structurally conserved molecules of microbes such as viruses, bacteria, and fungi and are an essential part of innate immunity (Netea et al. 2012). These functions of TLR genes could have led to balancing selective pressures on them.

However, no pLOF mutations were detected in TLR genes, the only gene family lacking such variants, indicating purifying selection. Several studies have found purifying selection acting on TLR genes (Stefanović et al. 2020; Mukherjee et al. 2014; Netea et al. 2012; Raven et al. 2017). These seemingly contradictory results - high heterozygosity alongside a lack of highly deleterious mutations - could reflect heterogeneous selection across the TLR genes, with purifying selection removing deleterious variants in some regions, while other regions are subject to selection favoring increased diversity. (Babik et al. 2015; Podlaszczuk et al. 2020). Nevertheless, both pieces of evidence point to potential mechanisms of genomic resilience, including purging and retention of functional diversity, that could contribute to population persistence. This analysis was restricted to SNP-derived loss-of-function mutations, with frameshift mutations excluded. As frameshift variants may contribute significantly to immune gene diversity, future investigations should extend this approach to include indel-based mutations.

We have not used traditional selection tests (such as Tajima’s D and McDonald-Kreitman’s test) to distinguish between neutral and non-neutral evolution, as these approaches do not distinguish between selective and demographic events (Poh et al. 2014). Given the drastic bottlenecks tigers have experienced throughout their evolutionary history (Armstrong et al. 2021), such tests would be uninformative. A previous study using the same Indian tiger reference found that some interleukin and chemokine genes were under positive selection, as determined by a phylogenetic approach (Shukla et al. 2023). Interestingly, all of the genes detected to be under positive selection were signalling genes, and none of the immune receptor genes were detected to be under positive selection. Our findings of balancing selection at receptor loci may help explain this pattern - balancing selection maintains diversity rather than fixing advantageous alleles, and would therefore not be detected by methods designed to identify positive directional selection. Together, these results suggest that receptor and signalling genes may be subject to fundamentally different selective pressures in the immune system.

### Mutation load in immune genes is similar to the genome-wide mutation load in inbred tiger populations

Estimating mutation load may help characterize variation in immune gene loci. Overall, load at immune genes is low, as only 30 potentially damaging loci were identified. Most of the missense variants were benign (Supplementary Figure 3). There were several missense mutations for which a Polyphen assignment was not possible, likely reflecting limitations associated with analyzing a non-model organism (Kovalev et al. 2018).

We found that the variation in damaging mutation load at immune genes met expextations of load in small, isolated versus large, connected populations (Khan et al., 2021). However, pLoF load was higher than pDeleterious load in the Northwest Indian tigers. We also observed higher pLoF than pDeleterious in South China tigers. Khan et al. 2021 found lower pLoF than pDeleterious in the Northwest Indian tiger populations and attributed it to purging. However, they did not investigate the possible pathogenicity of these mutations, which might explain the differences in results. Such damaging mutation load could be detrimental to the population when exposed to new pathogens (Asano et al. 2022).

### Mutation load and diversity shed light on populations of conservation importance

The genetic diversity of immune genes is known to facilitate the production of various protein motifs that can aid in identifying novel pathogens and impart disease resistance (Trowsdale and Parham 2004). Hence, loss of protective polymorphisms could lead to higher susceptibility to novel infectious diseases, potentially contributing to further population decline (Sommer 2005). Tasmanian devils are particularly notorious for their compromised immune function and increased susceptibility to facial tumors, which have contributed to a decline in their population (Morris et al. 2015; Strickland et al. 2024). Galapagos sea lions (Brock et al. 2015) and inbred song sparrows are other examples of inbred individuals exhibiting lowered immunity to parasitic infections (Reid et al. 2007).

Host immunity is a crucial factor for the survival of tigers and other carnivore populations in the future. Several species of large carnivores have suffered disease epidemics in the recent past, leading to population decline. Canine distemper virus epidemics have affected African wild dogs, African and Asiatic lions and Amur tigers (Mccarthy et al. 2007; Roelke-Parker et al. 2010; Gilbert et al. 2014; Mourya et al. 2025). Inbreeding in lions in South Africa is known to increase susceptibility to bovine tuberculosis.

In our study, we find that the Sumatran and South China subspecies exhibit low nucleotide diversity and heterozygosity, respectively, and are therefore most likely to be susceptible to new infections. The inbred Northwestern subpopulation of Indian tigers is also a population of concern, particularly given the prevalence of Canine Distemper Virus in the region (Sidhu et al. 2019). The reduced homozygous load and remnants of heterozygosity at certain immune genes in Northwest Indian and South China populations might yet help the populations persist even in the event of disease exposure. However, in the presence of genetic drift, this variation could eventually be lost (Gargiulo et al. 2024). Conservation measures, such as genetic rescue, must be cautious about the possibility of introducing deleterious alleles that are purged out in these inbred populations but are present as masked load in larger populations. However, we currently lack the experimental and epidemiological data needed to determine what range of immune gene variation is sufficient to mount an effective response to specific pathogens, or how variation at the loci studied here affects disease susceptibility or resistance in wild tiger populations. Addressing this gap will require integration of genomic data with longitudinal health monitoring and immunological assays, and represents an important direction for future conservation genomics research.

### Use and opportunities of genomics in studying adaptive variation of threatened species

Our study demonstrates the potential of utilizing genomic data beyond its initial research purpose, particularly for conservation, as we have not generated any new data and instead leveraged publicly available genomes. While our study highlights concordance between neutral and immunogenetic variation, the generality of such patterns warrants further investigation in other endangered species. As shown in Kardos et al. (2021), population genetic theory and empirical evidence from several studies indicate that neutral variation is correctly indicative of extinction risk, and genome-wide variation is important for population viability. A comprehensive understanding of functional variation can be obtained only for species with genomic information and access to samples, which is often limited to a few charismatic, endangered species. Our results contribute to the existing literature, which suggests that when such information is not available, genome-wide neutral markers can be indicative of adaptive potential. At the same time, previous studies have shown that adaptive loci may offer more direct insights for conservation planning in some cases (Oliver & Piertney 2012), and that gene-targeted approaches are particularly valuable when local adaptation or specific functional variants drive population persistence (Flanagan et al. 2018; Kardos and Shafer 2018). Together, these findings highlight the importance of integrating both neutral and genomic information,when available, to optimise conservation decision-making.

## Conclusion

Our study addresses an important but often overlooked aspect of tiger conservation genetics. We demonstrate that immunogenetic variation is correlated with neutral variation. Immune genes coding for receptor molecules have higher genetic variation than neutral loci in tigers. Populations that have undergone bottlenecks, inbreeding and founder effects exhibit lower immune gene (as well as neutral) diversity, implying a dominant role for neutral processes in such populations. However, despite drift, some receptor-coding immune genes and immune signaling loci retain high levels of polymorphism. Of these, TLR genes were found to exhibit a possible signature of balancing selection. We identified populations and subspecies with putatively disadvantageous variations in immune genes. Finally, in contexts where large-scale genomic data and information about functional variation is not available, our results suggest that neutral markers could adequately inform managers about the vulnerability of populations and species to extinction risks from pathogens and infectious diseases.

## Methods

### Identification of immune genes

Immune gene sequences were extracted from the annotation file of a female wild Indian Tiger genome assembly (GenBank ID: GCA_021130815.1) (PanTigT.MC.v3, Shukla et al. 2023). The associated General Feature Format file was examined to extract rows containing terms related to the immune genes of interest. We searched for genes coding for Toll-like receptors (TLRs), Leukocyte Immunoglobulin-Like receptors (LILRs), interleukins (ILs), chemokines (CC or CXC), and Tumor Necrosis Factor (TNF). These genes are well characterized and known to be involved in immunity (see Introduction). The search results were manually reviewed, and sequences with verified gene names and completely assembled Open Reading Frames were selected for further analysis. Genes were chosen for the study if the translated protein was an immune receptor or signaling molecule, and were excluded if they were present on a sex chromosome or on unplaced scaffolds. In cases where there were numerous entries of the same gene with similar start and end positions, the longest sequence was retained. The annotation also provided information on the start and end positions of genes and it’s exons on the PanTigT.MC.v3 reference. This allowed us to determine the start and end positions of the entire gene as well as the exons.

### Whole genome data reassembly

We downloaded and resequenced genomes of six subspecies of tigers from the European Nucleotide Archive and the China National GenBank in the form of raw FastQ files (Liu et al. 2018; Khan et al. 2021; Armstrong et al. 2021; Du et al. 2022; Zhang et al. 2023). Since data were derived from multiple sequencing projects, we only used whole genomes with a mean average depth of 10, and all raw reads were reprocessed through a unified pipeline, including identical trimming, mapping, and variant calling steps. In total, 107 genomes were downloaded, trimmed using the default parameters of Trimmomatic (Bolger et al. 2014) and adapter sequences as mentioned in the published studies, and then aligned to the same Indian tiger reference assembly PanTigT.MC.v3 used for gene annotation above, published in Shukla et al. (2023), using bwa-mem (Li 2013). The dataset includes genomes of both wild and captive individuals. Table 2 summarizes the sub species and population level sample sizes and more details of sample origin and accession numbers are listed in Supplementary Table 1.

**Table 2:** Sample sizes for each population and corresponding abbreviations used further in the text.

| Subspecies | Number of genomes |
| --- | --- |
| Amur (AMU) | 18 |
| Malayan (MAL) | 14 |
| Sumatran (SUM) | 7 |
| Indochinese (INC) | 6 |
| Bengal (BEN) | 43 |
| South China (SOC) | 19 |
| Indian subpopulations (part of the 43 Bengal genomes) |  |
| Northwest India (NW) | 19 |
| South India (SI) | 10 |
| Central India (CI) | 8 |

### SNP calling and filtering for neutral and immune loci

Since intergenic loci are not expected to be subject to selection, they were chosen as surrogates for neutral loci. We defined intergenic loci as regions 50 kb upstream and downstream of all genic regions throughout the genome, as described in Armstrong et al. (2021). The Variant Call Format (VCF) files, containing both variant and invariant sites, were created using bcftools (Danecek et al. 2011). An mpileup was created, followed by identification of variants which were annotated with the relevant depth tags using the --annotate tag. Using bcftools norm, redundant lines were eliminated, and variants were left aligned. VCFtools (Danecek et al., 2011) was used to filter the resulting two VCF files for standard quality filters, including a minimum base quality of 30 and a minimum genotype quality of 30, and to remove alleles with a minor allele count of three or less. We also exclude insertions and deletions since the identification of structural variants is not reliable with the SNP calling method we employ. Further, variant and invariant sites were subjected to the same filters. We excluded sites with a Hardy-Weinberg p-value less than 0.0001 (to account for genotyping errors) and retained only biallelic sites (to minimize errors). Sites that were present in fewer than 70% of the individuals were filtered out. Finally, sites with mean depth across individuals below the 2.5th percentile and above the 97.5th percentile were removed, as described in Khan et al. (2021), ensuring that high-coverage sites are retained while removing sites with exceptionally high coverage that might be repeats. Finally, the filtered variant and invariant sites VCFs were merged using bcftools concat command (Danecek et al. 2011).

### Nucleotide diversity, Heterozygosity, and F_ST_ estimation

Nucleotide diversity and heterozygosity were estimated in two ways for immune receptor, immune signalling, and intergenic loci: a) with all samples pooled as a single population, b) samples subset for each subspecies and each Indian subpopulation. The first approach aims to understand the distribution of diversity values for the immune receptor, immune signaling, and neutral regions in the tiger species. The second approach allows us to make the same comparison at the subspecies and subpopulation levels. Since South China tigers are completely captive and extinct in the wild, we excluded the South China (SOC) samples in the pooled analyses a) and include it in analyses b) where we highlight this population separately. The VCF containing exonic and intergenic variants was used as input for Pixy (Korunes and Samuk 2021) to estimate nucleotide diversity (π) and F_ST_ for non-overlapping 500 bp windows (the average size of exons is 391 bases, which we rounded off to arrive at the window size). Heterozygosity was calculated as the proportion of observed heterozygotes for every exonic site in immune genes and all sites in intergenic loci. Specifically, the –hardy option in VCFtools (Danecek et al., 2011) was used to obtain observed genotypes, and the proportion of observed heterozygotes at each SNP was calculated by summing the number of observed heterozygotes and dividing by the total number of observed genotypes. Pairwise Hudson’s F_ST_ (Hudson et al. 1992) was estimated for both immune and intergenic datasets to account for unequal sample size across different groupings. These distributions were visualized using boxplots. The subsets of pairwise F_ST_ were further visualized as follows: i) F_ST_ of Amur with all other subspecies, since Amur tigers were exposed to Canine Distemper Virus; ii) F_ST_ of Sumatran tigers with all other subspecies, since they underwent a founding event and exist on an island, and iii) the Rest of the subspecies and the Indian subpopulation level comparisons.

Throughout study, we use “locus/loci” as a general term for genomic regions of interest, which vary in scale depending on the analysis as described above. All the diversity metrics were visualized by categorizing the immune loci as receptor or signalling loci. This allowed us to account for varying gene sizes and statistically compare these metrics across categories. However, the categorization does not aggregate the calculation of the metrics.

### Comparison of site frequency spectrum to infer selection

To test for selection at immune genes, we estimated the unfolded site frequency spectrum (SFS) across all 107 genomes for synonymous and nonsynonymous SNPs, and compared this with the unfolded SFS at neutral loci. To account for the difference in the number of neutral and immune sites examined, we divided each bin by the total number of SNPs for that category. SFS was estimated using the angsd -doSaf 1 and realSFS programs, and was polarized using a genomic alignment of closely related species - domestic cat, lion, and cheetah - as described in Khan et al. (2021). With this approach, the type of selection can be inferred as follows: i) Purifying selection if the SFS of non-synonymous loci has fewer rare alleles when compared to neutral and synonymous SFS, ii) Positive selection if the SFS of non-synonymous loci has more rare alleles when compared to neutral and synonymous SFS, and iii) Balancing selection if there are more alleles at intermediate frequencies in the non-synonymous SFS when compared to neutral and synonymous SFS (Nielsen 2005).

To detect synonymous and nonsynonymous variants, Ensembl Variant Effect Prediction, release 99 (McLaren et al. 2016) was used to predict the effect of variants in immune genes. VEP was supplied with the reference FASTA and corresponding GFF files derived from the assembly used for read mapping (PanTigT.MC.v3, Shukla et al., 2023), with the tags “- everything” and “- pick-allele”. Variants predicted to be synonymous and missense were used to create SFS for synonymous and nonsynonymous SFS, respectively.

### Statistical tests

Differences in nucleotide diversity of immune receptor, signalling, and neutral loci between populations were tested using a Zero Inflated Gamma Generalized Linear Model implemented through glmmTMB R package (Brooks et al. 2017) (available on GitHub as well as CRAN). Model fit was evaluated using p-value and the estimate.

Differences in heterozygosity of immune receptor, signalling, and intergenic loci, as well as differences in site frequency spectra of neutral, synonymous, and nonsynonymous SNPs, were tested using a Binomial Generalized Linear Model. These models were evaluated using p-value as well as pseudo-R-squared.

Differences in pairwise Hudson’s F_ST_ were estimated with a Gaussian model implemented through the glm() function in R, and model fit was assessed using p-value and pseudo-R-squared. High values of pseudo-R-squared indicated a good model fit; the estimate in the zero-inflated GLM was used to assess deviation from the null hypothesis, which was the diversity at neutral loci in this study.

The differences in heterozygosity between populations were tested using the Kruskal-Wallis p-value test in RStudio, which is appropriate for comparing more than two groups without assuming normality.

To identify genes with high nucleotide diversity, we identified windows of signalling immune genes with nucleotide diversity in the top two quantiles and receptor immune genes in the top quantile of this distribution. Genes with high heterozygosity were identified based on upper bound boxplot outliers i.e. Q3+(1.5*IQR) where Q3 is the 75th percentile of the data and IQR is the interquartile range.

### Identification of deleterious mutation and estimation of mutation load

Results from Ensembl Variant Effect Prediction (McLaren et al., 2016) were used to identify deleterious mutations in immune genes. Missense variants can alter the structure of the protein product, but this change may be benign or deleterious. Hence, the potential pathogenicity of the identified missense variants was predicted using PolyPhen2 (Adzhubei et al. 2013) with the HumVar2 protocol, which compares known pathogenic amino acid changes in humans to classify query mutations as benign, possibly damaging, or probably damaging. Possibly and probably damaging variants were considered putatively deleterious variants (pDeleterious). Variants that were classified as putatively Loss of Function (pLOF) were transcript_ablation, splice_donor_variant, splice_acceptor_variant, stop_gained, and splice_region_variant. pDeleterious variants were assumed to be mildly deleterious, while pLOF variants were considered to be strongly deleterious.

Since load is defined by the number of derived deleterious alleles, we inferred the ancestral state for each pDeleterious and pLOF allele from the majority allele present in the genomic alignment of domestic cats, lions, and cheetahs, as described in Khan et al. (2021). Sites at which ancestral alleles were missing were excluded from the estimation of mutation load, while derived alleles were retained. We estimated the mutation load for Indian subpopulations to contrast the mutation load between the small, isolated, and inbred Northwest Indian population and the two large, connected Indian populations (South Indian and Central Indian). We also compared the load in inbred South China tigers with that of Malayan tigers, which are geographically close to the South China tigers but are not inbred. Mutation load was estimated in terms of the number of alleles using VCFtools –hwe tag to find the number of loci that were homozygous and heterozygous for pDeleterious and pLOF allele. Homozygous load helped us to understand realized or inbreeding load, whereas heterozygous load estimated the masked load in a population.

## Supporting information

ResultTables

## Acknowledgments

We are thankful to Laura Bertola, Hareendra Baraiya and Anubhab Khan for comments on the manuscript; Megan Aylward, Mayuresh Gangal and Mihir Joshi for discussion and inputs, and Kritagnya Vadar for help with mining annotations. BVA was supported by the NCBS/Tata Institute of Fundamental Research (TIFR) graduate program.

## Data availability statement

No new data was generated for this study. Supplementary tables 1 and 2 are in excel sheet and rest of the supplementary information is in word document. Accession information for the data used is available in the Supplementary Table 1.

## References

Acevedo-Whitehouse, Karina, and Andrew A. Cunningham. 2006. “Is MHC Enough for Understanding Wildlife Immunogenetics?” Trends in Ecology and Evolution 21 (8): 433–38. 10.1016/j.tree.2006.05.010.

Adzhubei, Ivan, Daniel M. Jordan, and Shamil R. Sunyaev. 2013. “Predicting Functional Effect of Human Missense Mutations Using PolyPhen-2.” Current Protocols in Human Genetics 76 (1): 7.20.1-7.20.41. 10.1002/0471142905.hg0720s76.

Aguilar, Andres, Gary Roemer, Sally Debenham, Matthew Binns, David Garcelon, and Robert K. Wayne. 2004. “High MHC Diversity Maintained by Balancing Selection in an Otherwise Genetically Monomorphic Mammal.” Proceedings of the National Academy of Sciences of the United States of America 101 (10): 3490–94. 10.1073/PNAS.0306582101.

Almeida Lopes, Camila de, Linda Djune Yemeli, and Pedro Gazzinelli-Guimaraes. 2025. “Innate Immune Recognition of Helminths: TLRs and Beyond.” Parasite Immunology 47 (12): e70042. 10.1111/pim.70042.

Areal, Helena, Joana Abrantes, and Pedro J. Esteves. 2011. “Signatures of Positive Selection in Toll-like Receptor (TLR) Genes in Mammals.” BMC Evolutionary Biology 11 (1): 368. 10.1186/1471-2148-11-368.

Armstrong, Ellie E., Anubhab Khan, Ryan W. Taylor, et al. 2021. “Recent Evolutionary History of Tigers Highlights Contrasting Roles of Genetic Drift and Selection.” Molecular Biology and Evolution 38 (6): 2021. 10.1093/molbev/msab032.

Asano, Takaki, Takanori Utsumi, Reiko Kagawa, Shuhei Karakawa, and Satoshi Okada. 2022. “Inborn Errors of Immunity with Loss- and Gain-of-Function Germline Mutations in STAT1.” Clinical and Experimental Immunology 212 (2): 96–106. 10.1093/cei/uxac106.

Babik, Wieslaw, Katarzyna Dudek, Anna Fijarczyk, et al. 2015. “Constraint and Adaptation in Newt Toll-Like Receptor Genes.” Genome Biology and Evolution 7 (1): 81–95. 10.1093/GBE/EVU266.

Bernhardsson, Carolina, and Pär K. Ingvarsson. 2011. “Molecular Population Genetics of Elicitor-Induced Resistance Genes in European Aspen (Populus Tremula L., Salicaceae).” PLOS ONE 6 (9): e24867. 10.1371/journal.pone.0024867.

Bhattacharya, Sanchita, Patrick Dunn, Cristel G. Thomas, et al. 2018. “ImmPort, toward Repurposing of Open Access Immunological Assay Data for Translational and Clinical Research.” Scientific Data 5 (1): 180015. 10.1038/sdata.2018.15.

Bi, Jinping, Xiaoyun Hu, Dejun Mu, et al. 2026. “Gene Expression and Immune Cell Heterogeneity in Inbred Amur Tiger.” BMC Genomics, ahead of print, April 25. 10.1186/s12864-026-12872-y.

Bitarello, Bárbara D., Cesare de Filippo, João C. Teixeira, et al. 2018. “Signatures of Long-Term Balancing Selection in Human Genomes.” Genome Biology and Evolution 10 (3): 939– 55. 10.1093/gbe/evy054.

Bolger, Anthony M., Marc Lohse, and Bjoern Usadel. 2014. “Trimmomatic: A Flexible Trimmer for Illumina Sequence Data.” Bioinformatics 30 (15): 2114. 10.1093/BIOINFORMATICS/BTU170.

Bollmer, Jennifer L., Elizabeth A. Ruder, Jeff A. Johnson, John A. Eimes, and Peter O. Dunn. 2011. “Drift and Selection Influence Geographic Variation at Immune Loci of Prairie-Chickens.” Molecular Ecology 20 (22): 4695–706. 10.1111/J.1365-294X.2011.05319.X.

Bosse, Mirte, and Sam van Loon. 2022. “Challenges in Quantifying Genome Erosion for Conservation.” Frontiers in Genetics 13 (September). 10.3389/fgene.2022.960958.

Brandt, Débora Y. C., Jônatas César, Jérôme Goudet, and Diogo Meyer. 2018. “The Effect of Balancing Selection on Population Differentiation: A Study with HLA Genes.” G3: Genes, Genomes, Genetics 8 (8): 2805–15. 10.1534/g3.118.200367.

Brock, Patrick M., Simon J. Goodman, Ailsa J. Hall, Marilyn Cruz, and Karina Acevedo-whitehouse. 2015. “Context-Dependent Associations between Heterozygosity and Immune Variation in a Wild Carnivore.” BMC Evolutionary Biology, 1–10. 10.1186/s12862-015-0519-6.

Brooks, Mollie, E., Kasper Kristensen, Koen Benthem J., van, et al. 2017. “glmmTMB Balances Speed and Flexibility Among Packages for Zero-Inflated Generalized Linear Mixed Modeling.” The R Journal 9 (2): 378. 10.32614/RJ-2017-066.

Capozza, Paolo, Annamaria Pratelli, Michele Camero, et al. 2021. “Feline Coronavirus and Alpha-Herpesvirus Infections: Innate Immune Response and Immune Escape Mechanisms.” Animals 11 (12): 3548. 10.3390/ani11123548.

Castro-Prieto, Aines, Bettina Wachter, and Simone Sommer. 2010. “Cheetah Paradigm Revisited: MHC Diversity in the World’s Largest Free-Ranging Population.” Molecular Biology and Evolution 28 (4): 1455–68. 10.1093/molbev/msq330.

Clegg, Sonya M., Sandie M. Degnan, Craig Moritz, Arnaud Estoup, Jiro Kikkawa, and Ian P. F. Owens. 2002. “MICROEVOLUTION IN ISLAND FORMS: THE ROLES OF DRIFT AND DIRECTIONAL SELECTION IN MORPHOLOGICAL DIVERGENCE OF A PASSERINE BIRD.” Evolution 56 (10): 2090–99. 10.1111/j.0014-3820.2002.tb00134.x.

Cortazar-Chinarro, Maria, Sara Meurling, Laurens Schroyens, et al. 2022. “Major Histocompatibility Complex Variation and Haplotype Associated Survival in Response to Experimental Infection of Two Bd-GPL Strains Along a Latitudinal Gradient.” Frontiers in Ecology and Evolution 10 (July). 10.3389/fevo.2022.915271.

Danecek, Petr, Adam Auton, Goncalo Abecasis, et al. 2011. “The Variant Call Format and VCFtools.” Bioinformatics 27 (15): 2156–58. 10.1093/bioinformatics/btr330.

Davies, Charli S., Martin I. Taylor, Martijn Hammers, et al. 2021. “Contemporary Evolution of the Innate Immune Receptor Gene TLR3 in an Isolated Vertebrate Population.” Molecular Ecology 30 (11): 2528–42. 10.1111/mec.15914.

Du, Hairong, Jingjing Yu, Qian Li, and Minghai Zhang. 2022. “New Evidence of Tiger Subspecies Differentiation and Environmental Adaptation: Comparison of the Whole Genomes of the Amur Tiger and the South China Tiger.” Animals 12 (July): 1817. 10.3390/ani12141817.

Ebert, Dieter, and Peter D. Fields. 2020. “Host-Parasite Co-Evolution and Its Genomic Signature.” Nature Reviews. Genetics 21 (12): 754–68. 10.1038/s41576-020-0269-1.

Edwards, Scott V., Joe Gasper, Daniel Garrigan, Duane Martindale, and Ben F. Koop. 2000. “A 39-Kb Sequence around a Blackbird Mhc Class II Gene: Ghost of Selection Past and Songbird Genome Architecture.” Molecular Biology and Evolution 17 (9): 1384–95. 10.1093/oxfordjournals.molbev.a026421.

Ejsmond, Maciej Jan, and Jacek Radwan. 2015. “Red Queen Processes Drive Positive Selection on Major Histocompatibility Complex (MHC) Genes.” PLOS Computational Biology 11 (11): e1004627. 10.1371/journal.pcbi.1004627.

Fijarczyk, Anna, and Wiesław Babik. 2015. “Detecting Balancing Selection in Genomes: Limits and Prospects.” Molecular Ecology 24 (14): 3529–45. 10.1111/mec.13226.

Fijarczyk, Anna, Katarzyna Dudek, and Wieslaw Babik. 2016. “Selective Landscapes in Newt Immune Genes Inferred from Patterns of Nucleotide Variation.” Genome Biology and Evolution 8 (11): 3417–32. 10.1093/gbe/evw236.

Flanagan, Sarah P., Brenna R. Forester, Emily K. Latch, Sally N. Aitken, and Sean Hoban. 2018. “Guidelines for Planning Genomic Assessment and Monitoring of Locally Adaptive Variation to Inform Species Conservation.” Evolutionary Applications 11 (7): 1035–52. 10.1111/eva.12569.

Frankham, Richard. 1996. “Relationship of Genetic Variation to Population Size in Wildlife.” Conservation Biology 10 (6): 1500–1508. 10.1046/j.1523-1739.1996.10061500.x.

Gargiulo, Roberta, Katharina B. Budde, and Myriam Heuertz. 2024. “Mind the Lag: Understanding Genetic Extinction Debt for Conservation.” Trends in Ecology & Evolution, ahead of print, November 20. 10.1016/j.tree.2024.10.008.

Gilbert, Martin, Zachary Dvornicky-Raymond, and Jessica Bodgener. 2023a. “Disease Threats to Tigers and Their Prey.” Frontiers in Ecology and Evolution 11 (April). 10.3389/fevo.2023.1135935.

Gilbert, Martin, Zachary Dvornicky-Raymond, and Jessica Bodgener. 2023b. “Disease Threats to Tigers and Their Prey.” Frontiers in Ecology and Evolution 11 (April). 10.3389/fevo.2023.1135935.

Gilbert, Martin, Zachary Dvornicky-Raymond, and Jessica Bodgener. 2023c. “Disease Threats to Tigers and Their Prey.” Frontiers in Ecology and Evolution 11 (April): 1135935. 10.3389/fevo.2023.1135935.

Gilbert, Martin, Dale G. Miquelle, John M. Goodrich, et al. 2014. “Estimating the Potential Impact of Canine Distemper Virus on the Amur Tiger Population (Panthera Tigris Altaica) in Russia.” PLOS ONE 9 (10): e110811. 10.1371/JOURNAL.PONE.0110811.

Gilroy, D. L., K. P. Phillips, D. S. Richardson, and C. van Oosterhout. 2017. “Toll-like Receptor Variation in the Bottlenecked Population of the Seychelles Warbler: Computer Simulations See the ‘ghost of Selection Past’ and Quantify the ‘drift Debt’.” Journal of Evolutionary Biology 30 (7): 1276–87. 10.1111/jeb.13077.

Grossen, Christine, and Uma Ramakrishnan. 2024. “Genetic Load.” Current Biology 34 (24): R1216–20. 10.1016/j.cub.2024.11.004.

Haddad, Nick M., Lars A. Brudvig, Jean Clobert, et al. 2015. “Habitat Fragmentation and Its Lasting Impact on Earth’s Ecosystems.” Science Advances 1 (2): e1500052. 10.1126/sciadv.1500052.

Hasselgren, Malin, Nicolas Dussex, Johanna von Seth, Anders Angerbjörn, Love Dalén, and Karin Norén. 2024. “Strongly Deleterious Mutations Influence Reproductive Output and Longevity in an Endangered Population.” Nature Communications 15 (1): 8378. 10.1038/s41467-024-52741-4.

Hudson, R. R., M. Slatkin, and W. P. Maddison. 1992. “Estimation of Levels of Gene Flow from DNA Sequence Data.” Genetics 132 (2): 583–89. 10.1093/genetics/132.2.583.

Jhala, Yadvendradev V., Ninad Avinash Mungi, Rajesh Gopal, and Qamar Qureshi. 2025. “Tiger Recovery amid People and Poverty.” Science 387 (6733): 505–10. 10.1126/science.adk4827.

Kardos, Marty, and Aaron B. A. Shafer. 2018. “The Peril of Gene-Targeted Conservation.” Trends in Ecology & Evolution 33 (11): 827–39. 10.1016/j.tree.2018.08.011.

Keller, Lukas F., and Donald M. Waller. 2002. Inbreeding Effects in Wild Populations. 17 (5): 19–23.

Khan, Anubhab, Kaushalkumar Patel, Harsh Shukla, et al. 2021. “Genomic Evidence for Inbreeding Depression and Purging of Deleterious Genetic Variation in Indian Tigers.” Proceedings of the National Academy of Sciences 118 (49): e2023018118. 10.1073/PNAS.2023018118.

Kloch, Agnieszka, Marius A. Wenzel, Dominik R. Laetsch, Olek Michalski, Renata Welc-Fall1ciak, and Stuart B. Piertney. 2018. “Signatures of Balancing Selection in Toll-like Receptor (TLRs) Genes - Novel Insights from a Free-Living Rodent.” Scientific Reports 8 (1): 1–10. 10.1038/s41598-018-26672-2.

Korunes, Katharine L., and Kieran Samuk. 2021. “Pixy: Unbiased Estimation of Nucleotide Diversity and Divergence in the Presence of Missing Data.” Molecular Ecology Resources 21 (4): 1359–68. 10.1111/1755-0998.13326.

Kovalev, Maxim S., Anna A. Igolkina, Maria G. Samsonova, and Sergey V. Nuzhdin. 2018. “A Pipeline for Classifying Deleterious Coding Mutations in Agricultural Plants.” Frontiers in Plant Science 9 (November). 10.3389/fpls.2018.01734.

Lara, Carlos Esteban, Catherine E. Grueber, Benedikt Holtmann, et al. 2020. “Assessment of the Dunnocks’ Introduction to New Zealand Using Innate Immune-Gene Diversity.” Evolutionary Ecology 34 (5): 803–20. 10.1007/s10682-020-10070-0.

Li, Heng. 2013. “Aligning Sequence Reads, Clone Sequences and Assembly Contigs with BWA-MEM.” arXiv:1303.3997. Preprint, arXiv, May 26. 10.48550/arXiv.1303.3997.

Liu, Yue Chen, Xin Sun, Carlos Driscoll, et al. 2018. “Genome-Wide Evolutionary Analysis of Natural History and Adaptation in the World’s Tigers.” Current Biology 28 (23): 3840–3849.e6. 10.1016/j.cub.2018.09.019.

Luo, Mao Fang, Hui Juan Pan, Zhi Jin Liu, and Ming Li. 2012. “Balancing Selection and Genetic Drift at Major Histocompatibility Complex Class II Genes in Isolated Populations of Golden Snub-Nosed Monkey (Rhinopithecus Roxellana).” BMC Evolutionary Biology 12 (1): 207. 10.1186/1471-2148-12-207.

Mable, Barbara K. 2019. “Conservation of Adaptive Potential and Functional Diversity: Integrating Old and New Approaches.” In Conservation Genetics, vol. 20. no. 1. Springer Netherlands, February. 10.1007/s10592-018-1129-9.

“Major Histocompatibility Complex Class I Polymorphism in Asiatic Lions - Sachdev - 2005 - Tissue Antigens - Wiley Online Library.” 2025. January 15. https://onlinelibrary.wiley.com/doi/full/10.1111/j.1399-0039.2005.00432.x.

Martin, Katherine R., Katherine L. Mansfield, and Anna E. Savage. 2022. “Adaptive Evolution of Major Histocompatibility Complex Class I Immune Genes and Disease Associations in Coastal Juvenile Sea Turtles.” Royal Society Open Science 9 (2): 211190. 10.1098/rsos.211190.

Mathur, Samarth, Andrew J. Mason, Gideon S. Bradburd, and H. Lisle Gibbs. 2023. “Functional Genomic Diversity Is Correlated with Neutral Genomic Diversity in Populations of an Endangered Rattlesnake.” Proceedings of the National Academy of Sciences of the United States of America 120 (43). 10.1073/pnas.2303043120.

Mccarthy, Alex J., Marie-anne Shaw, and Simon J. Goodman. 2007. Pathogen Evolution and Disease Emergence in Carnivores. no. October: 3165–74. 10.1098/rspb.2007.0884.

McCauley, Deborah, Virginia Stout, Kamal P. Gairhe, et al. 2021. “Serologic Survey of Selected Pathogens in Free-Ranging Bengal Tigers (Panthera Tigris Tigris) in Nepal.” Journal of Wildlife Diseases 57 (2). 10.7589/JWD-D-20-00046.

McLaren, William, Laurent Gil, Sarah E. Hunt, et al. 2016. “The Ensembl Variant Effect Predictor.” Genome Biology 17 (1): 122. 10.1186/s13059-016-0974-4.

Mondol, Samrat, Michael W. Bruford, and Uma Ramakrishnan. 2013. Demographic Loss, Genetic Structure and the Conservation Implications for Indian Tigers.

Morris, Katrina M., Belinda Wright, Catherine E. Grueber, Carolyn Hogg, and Katherine Belov. 2015. “Lack of Genetic Diversity across Diverse Immune Genes in an Endangered Mammal, the Tasmanian Devil (Sarcophilus Harrisii).” Molecular Ecology 24 (15): 3860–72. 10.1111/mec.13291.

Mourya, Devendra T., Pragya D. Yadav, Sreelekshmy Mohandas, et al. 2025. Canine Distemper Virus in Asiatic Lions of Gujarat State, India - Volume 25, Number 11—November 2019 - Emerging Infectious Diseases Journal - CDC. February 10. 10.3201/eid2511.190120.

Mukherjee, Souvik, Debdutta Ganguli, and Partha P. Majumder. 2014. “Global Footprints of Purifying Selection on Toll-Like Receptor Genes Primarily Associated with Response to Bacterial Infections in Humans.” Genome Biology and Evolution 6 (3): 551–58. 10.1093/gbe/evu032.

Mukherjee, Souvik, Neeta Sarkar-Roy, Diane K. Wagener, and Partha P. Majumder. 2009. “Signatures of Natural Selection Are Not Uniform across Genes of Innate Immune System, but Purifying Selection Is the Dominant Signature.” Proceedings of the National Academy of Sciences 106 (17): 7073–78. 10.1073/PNAS.0811357106.

Natesh, Meghana, Goutham Atla, Parag Nigam, et al. 2017. “Conservation Priorities for Endangered Indian Tigers through a Genomic Lens.” Scientific Reports 7 (1): 1–11. 10.1038/s41598-017-09748-3.

Netea, Mihai G., Cisca Wijmenga, and Luke A. J. O’Neill. 2012. “Genetic Variation in Toll-like Receptors and Disease Susceptibility.” Nature Immunology 13 (6): 535–42. 10.1038/ni.2284.

Nielsen, Rasmus. 2005. “Molecular Signatures of Natural Selection.” Annual Review of Genetics 39 (Volume 39, 2005): 197–218. 10.1146/annurev.genet.39.073003.112420.

Oliver, Matthew K., and Stuart B. Piertney. 2012. “Selection Maintains MHC Diversity through a Natural Population Bottleneck.” Molecular Biology and Evolution 29 (7): 1713–20. 10.1093/molbev/mss063.

Podlaszczuk, Patrycja, Piotr Indykiewicz, Janusz Markowski, and Piotr Minias. 2020. “Relaxation of Selective Constraints Shapes Variation of Toll-like Receptors in a Colonial Waterbird, the Black-Headed Gull.” Immunogenetics 72 (4): 251. 10.1007/S00251-020-01156-8.

Poh, Yu Ping, Vera S. Domingues, Hopi E. Hoekstra, and Jeffrey D. Jensen. 2014. “On the Prospect of Identifying Adaptive Loci in Recently Bottlenecked Populations.” PLoS ONE 9 (11). 10.1371/journal.pone.0110579.

Pokorny, Ina, Reeta Sharma, and Ralph Tiedemann. 2010. MHC Class I and MHC Class II DRB Gene Variability in Wild and Captive Bengal Tigers (Panthera Tigris Tigris). 667–79. 10.1007/s00251-010-0475-7.

Raven, Nynke, Simeon Lisovski, Marcel Klaassen, et al. 2017. “Purifying Selection and Concerted Evolution of RNA-Sensing Toll-like Receptors in Migratory Waders.” Infection, Genetics and Evolution 53 (September): 135–45. 10.1016/j.meegid.2017.05.012.

Reid, Jane M., Peter Arcese, Lukas F. Keller, Kyle H. Elliott, Laura Sampson, and Dennis Hasselquist. 2007. Inbreeding Effects on Immune Response in Free-Living Song Sparrows (Melospiza Melodia). no. November 2006: 697–706. 10.1098/rspb.2006.0092.

Roelke-Parker, M. E., L. Munson, C. Packer, et al. 2010. “A Canine Distemper Virus Epidemic in Serengeti Lions (Panthera Leo) (Vol 379, Pg 441, 1996).” Nature 464 (7290): 942. Doi%2010.1038/Nature08888.

S, Sommer. 2005. “The Importance of Immune Gene Variability (MHC) in Evolutionary Ecology and Conservation.” Frontiers in Zoology 2 (October). 10.1186/1742-9994-2-16.

Schubert, Nadine, Hazel J. Nichols, Francis Mwanguhya, et al. 2025. “Inbreeding Does Not Reduce Major Histocompatibility Complex Diversity in the Banded Mongoose.” BMC Ecology and Evolution 25 (1): 104. 10.1186/s12862-025-02456-x.

Shukla, Harsh, Kushal Suryamohan, Anubhab Khan, et al. 2023. “Near-Chromosomal de Novo Assembly of Bengal Tiger Genome Reveals Genetic Hallmarks of Apex Predation.” GigaScience 12. 10.1093/gigascience/giac112.

Sidhu, Nadisha, Jimmy Borah, Sunny Shah, Nidhi Rajput, and Kajal Kumar Jadav. 2019. “Is Canine Distemper Virus (CDV) a Lurking Threat to Large Carnivores? A Case Study from Ranthambhore Landscape in Rajasthan, India.” Journal of Threatened Taxa 11 (9): 14220–23. 10.11609/jott.4569.11.9.14220-14223.

Silver, Luke W., Elspeth A. McLennan, Julian Beaman, et al. 2024. “Using Bioinformatics to Investigate Functional Diversity: A Case Study of MHC Diversity in Koalas.” Immunogenetics 76 (5): 381–95. 10.1007/s00251-024-01356-6.

Soni, Vivak, and Jeffrey D. Jensen. 2024. “Temporal Challenges in Detecting Balancing Selection from Population Genomic Data.” G3 Genes|Genomes|Genetics 14 (6): jkae069. 10.1093/g3journal/jkae069.

Stefanović, Milomir, Mihajla Djan, Nevena Veličković, et al. 2020. “Purifying Selection Shaping the Evolution of the Toll-like Receptor 2 TIR Domain in Brown Hares (Lepus Europaeus) from Europe and the Middle East.” Molecular Biology Reports 47 (4): 2975–84. 10.1007/s11033-020-05382-x.

Strand, Tanja M., Gernot Segelbacher, María Quintela, Lingyun Xiao, Tomas Axelsson, and Jacob Höglund. 2012a. “Can Balancing Selection on MHC Loci Counteract Genetic Drift in Small Fragmented Populations of Black Grouse?” Ecology and Evolution 2 (2): 341–53. 10.1002/ece3.86.

Strand, Tanja M., Gernot Segelbacher, María Quintela, Lingyun Xiao, Tomas Axelsson, and Jacob Höglund. 2012b. “Can Balancing Selection on MHC Loci Counteract Genetic Drift in Small Fragmented Populations of Black Grouse?” Ecology and Evolution 2 (2): 341–53. 10.1002/ece3.86.

Strickland, Kasha, Menna E. Jones, Andrew Storfer, et al. 2024. “Adaptive Potential in the Face of a Transmissible Cancer in Tasmanian Devils.” Molecular Ecology 33 (21): e17531. 10.1111/mec.17531.

Trowsdale, John, and Peter Parham. 2004. “Defense Strategies and Immunity-Related Genes.” In European Journal of Immunology, vol. 34. no. 1. Eur J Immunol, January. 10.1002/eji.200324693.

Turner, Andrew K., Mike Begon, Joseph A. Jackson, Janette E. Bradley, and Steve Paterson. 2011. “Genetic Diversity in Cytokines Associated with Immune Variation and Resistance to Multiple Pathogens in a Natural Rodent Population.” PLoS Genetics 7 (10): e1002343. 10.1371/journal.pgen.1002343.

Vlček, Jakub, Matěj Miláček, Michal Vinkler, and Jan Štefka. 2023. “Effect of Population Size and Selection on Toll-like Receptor Diversity in Populations of Galápagos Mockingbirds.” Journal of Evolutionary Biology 36 (1): 109–20. 10.1111/JEB.14121.

Weedall, Gareth D., and David J. Conway. 2010. “Detecting Signatures of Balancing Selection to Identify Targets of Anti-Parasite Immunity.” Trends in Parasitology 26 (7): 363–69. 10.1016/j.pt.2010.04.002.

Wei, Kun, Zhihe Zhang, Xiaofang Wang, et al. 2010. “Lineage Pattern, Trans-Species Polymorphism, and Selection Pressure among the Major Lineages of Feline Mhc-DRB Peptide-Binding Region.” Immunogenetics 62 (5): 307–17. 10.1007/s00251-010-0440-5.

Willi, Yvonne, Torsten N. Kristensen, Carla M. Sgrò, Andrew R. Weeks, Michael Ørsted, and Ary A. Hoffmann. 2022. “Conservation Genetics as a Management Tool: The Five Best-Supported Paradigms to Assist the Management of Threatened Species.” Proceedings of the National Academy of Sciences 119 (1): e2105076119. 10.1073/pnas.2105076119.

Zhang, Le, Tianming Lan, Chuyu Lin, et al. 2023. “Chromosome-Scale Genomes Reveal Genomic Consequences of Inbreeding in the South China Tiger: A Comparative Study with the Amur Tiger.” Molecular Ecology Resources 23 (2): 330–47. 10.1111/1755-0998.13669.

