## Supplementary material for "Understanding range-wide immune gene variation in an endangered big cat (*Panthera tigris*)": ResultTables

**Supplementary Table 3: Mean (with 95 % confidence intervals) for nucleotide diversity and heterozygosity values at each gene family**

| **Gene Family** | **Mean** | **CI_L** | **CI_U** |
| --- | --- | --- | --- |
| **Nucleotide diversity** | | | |
| **TLR** | 0.0021 | 0.0005 | 0.0054 |
| **LILR** | 0.0012 | 0.0003 | 0.0029 |
| **Chemokine** | 0.0004 | 0.0003 | 0.0005 |
| **TNF** | 0.0004 | 0.0003 | 0.0005 |
| **Interleukin** | 0.0002 | 0.0001 | 0.0004 |
| **Heterozygosity** | | | |
| **TLR** | 0.253 | 0.197 | 0.308 |
| **LILR** | 0.119 | 0.086 | 0.155 |
| **Chemokine** | 0.157 | 0.134 | 0.180 |
| **TNF** | 0.165 | 0.146 | 0.187 |
| **Interleukin** | 0.175 | 0.134 | 0.221 |

**Supplementary Table 4: Zero-inflated Gamma Generalized Linear models for nucleotide diversity differences at family of loci**

| **Type of loci** | **Estimate** | **p-value** |
| --- | --- | --- |
| **Neutral** | 806.7703 |  |
| **Chemokine** | -6.8406 | 0.919 |
| **Interleukin** | 39.3557 | 0.797 |
| **LILR** | -542.5585 | <2e-16*** |
| **TLR** | -584.584 | <2e-16*** |
| **TNF** | -6.0739 | 0.899 |

**Supplementary Table 5: Binomial Generalized Linear models for heterozygosity differences at different family of loci**

| **Comparison vs neutral loci** | **p-value** | **Pseudo-R-squared** |
| --- | --- | --- |
| **TLR** | 0.4048 | 1.60E-05 |
| **LILR** | 0.0911 |  |
| **Interleukin** | 0.6775 |  |
| **TNF** | 0.1964 |  |
| **Chemokine** | 0.2112 |  |

**Supplementary Table 6: Mean (with 95 percent confidence intervals) of nucleotide diversity for each tiger subspecies and subpopulation**

| **Pop** | **Class** | **Mean** | **CI_L** | **CI_U** |
| --- | --- | --- | --- | --- |
| **Tiger Subspecies** | | | | |
| **SOC** | **Receptor** | 0.003009 | 0.000676 | 0.0072 |
| **MAL** | **Receptor** | 0.003477 | 0.000471 | 0.00824 |
| **INC** | **Receptor** | 0.000514 | 0.000285 | 0.000793 |
| **SUM** | **Receptor** | 0.001076 | 0.000174 | 0.002741 |
| **BEN** | **Receptor** | 0.001041 | 0.000475 | 0.001767 |
| **AMU** | **Receptor** | 0.002142 | 0.000349 | 0.005663 |
| **AMU** | **Signalling** | 0.000247 | 0.000177 | 0.000329 |
| **SUM** | **Signalling** | 0.000183 | 0.000126 | 0.000243 |
| **BEN** | **Signalling** | 0.000371 | 0.000287 | 0.000462 |
| **INC** | **Signalling** | 0.000338 | 0.000243 | 0.000436 |
| **SOC** | **Signalling** | 0.000369 | 0.000267 | 0.000481 |
| **MAL** | **Signalling** | 0.000331 | 0.000244 | 0.000433 |
| **BEN** | **Neutral** | 0.000641 | 0.000644 | 0.000638 |
| **MAL** | **Neutral** | 0.000775 | 0.000772 | 0.000772 |
| **SUM** | **Neutral** | 0.000392 | 0.000395 | 0.00039 |
| **INC** | **Neutral** | 0.000831 | 0.000839 | 0.000824 |
| **SOC** | **Neutral** | 0.000755 | 0.000758 | 0.000752 |
| **AMU** | **Neutral** | 0.000722 | 0.000725 | 0.000719 |
| **Indian Subpopulations** | | | | |
| **SI** | **Receptor** | 0.001241 | 0.000306 | 0.002702 |
| **CI** | **Receptor** | 0.001272 | 0.000453 | 0.00245 |
| **NW** | **Receptor** | 0.000403 | 0.000195 | 0.000656 |
| **SI** | **Signalling** | 0.000354 | 0.000259 | 0.000467 |
| **CI** | **Signalling** | 0.000392 | 0.000259 | 0.000467 |
| **NW** | **Signalling** | 0.000256 | 0.000177 | 0.000343 |
| **SI** | **Neutral** | 0.000564 | 0.000561 | 0.000567 |
| **CI** | **Neutral** | 0.000649 | 0.000646 | 0.000652 |
| **NW** | **Neutral** | 0.000467 | 0.000465 | 0.000465 |

**Supplementary Table 7: Mean (with 95 percent confidence intervals) of heterozygosity for each tiger subspecies and subpopulation**

| **Pop** | **Class** | **Mean** | **CI_L** | **CI_U** |
| --- | --- | --- | --- | --- |
| **Tiger Subspecies** | | | | |
| **INC** | **Neutral** | 0.2699933 | 0.269687 | 0.270287 |
| **INC** | **Immune** | 0.2380631 | 0.208323 | 0.269377 |
| **SOC** | **Neutral** | 0.1578823 | 0.157635 | 0.158125 |
| **SOC** | **Immune** | 0.1202035 | 0.103682 | 0.136994 |
| **BEN** | **Immune** | 0.1546989 | 0.138728 | 0.171292 |
| **BEN** | **Neutral** | 0.2068901 | 0.206663 | 0.207118 |
| **AMU** | **Immune** | 0.1418389 | 0.121969 | 0.162703 |
| **AMU** | **Neutral** | 0.1944994 | 0.194261 | 0.194728 |
| **MAL** | **Immune** | 0.2143697 | 0.192146 | 0.237049 |
| **MAL** | **Neutral** | 0.2485687 | 0.248297 | 0.248852 |
| **SUM** | **Neutral** | 0.1692154 | 0.168894 | 0.169512 |
| **SUM** | **Immune** | 0.1817819 | 0.153117 | 0.212182 |
| **Indian Subpopulations** | | | | |
| **NW** | **Immune** | 0.1219376 | 0.104002 | 0.142584 |
| **NW** | **Neutral** | 0.1559833 | 0.155712 | 0.156272 |
| **SI** | **Immune** | 0.1364535 | 0.116812 | 0.157006 |
| **SI** | **Neutral** | 0.2180707 | 0.217778 | 0.218386 |
| **CI** | **Immune** | 0.2182638 | 0.192578 | 0.245592 |
| **CI** | **Neutral** | 0.2675944 | 0.267254 | 0.267932 |

**Supplementary Table 8: Zero inflated Gamma models for testing differences in nucleotide diversity values for each subspecies**

| **Population/Type of loci** | **Estimate** | **p-value** |
| --- | --- | --- |
| **SUM** |  |  |
| **Neutral** | 665.6042 |  |
| **Receptor** | -513.3097 | <2e-16 *** |
| **Signalling** | -56.8677 | 0.36 |
| **SOC** |  |  |
| **Neutral** | 462.1204 | < 2e-16 *** |
| **Receptor** | -368.7958 | < 2e-16 *** |
| **Signalling** | -111.5752 | 0.000273 *** |
| **MAL** |  |  |
| **Neutral** | -6.352689 |  |
| **Receptor** | 1.987023 | < 2e-16 *** |
| **Signalling** | 0.165251 | 0.0351 * |
| **INC** |  |  |
| **Neutral** | 546.5976 |  |
| **Receptor** | -80.6352 | 0.2041 |
| **Signalling** | -91.978 | 0.0117 * |
| **BEN** |  |  |
| **Neutral** | 805.3779 |  |
| **Receptor** | -482.9071 | < 2e-16 *** |
| **Signalling** | -228.4885 | -5.1 3.12e-07 *** |
| **AMU** |  |  |
| **Neutral** | 648.9781 |  |
| **Receptor** | -533.8998 | < 2e-16 *** |
| **Signalling** | 25.5034 | 0.649 |

**Supplementary Table 9: Zero inflated Gamma models for testing differences in nucleotide diversity values for each Indian population**

| **Population/Type of loci** | **Estimate** | **p-value** |
| --- | --- | --- |
| **SI** |  |  |
| **Neutral** | 657.2025 |  |
| **Receptor** | -475.0462 | < 2e-16 *** |
| **Signalling** | -283.6099 | < 2e-16 *** |
| **CI** |  |  |
| **Neutral** | 613.6565 |  |
| **Receptor** | -398.3336 | < 2e-16 *** |
| **Signalling** | -183.1784 | 3.08e-08 *** |
| **NW** |  |  |
| **Neutral** | 588.4412 |  |
| **Receptor** | -163.6637 | 0.0154* |
| **Signalling** | -231.5707 | 2.64E-10 |

**Supplementary Table 10: Binomial models for testing differences in heterozygosity values of immune loci vs neutral loci in each subspecies**

| **Population** | **p-value** | **Pseudo-R-squared** |
| --- | --- | --- |
| **Tiger sub-species** |  |  |
| **SUM** | 0.481 | 0.00E+00 |
| **MAL** | 0.0956 | 2.27E-06 |
| **AMU** | 0.00528 | 7.33E-06 |
| **INC** | 0.13 | 1.89E-06 |
| **SOC** | 0.0299 | 4.24E-06 |
| **BEN** | 0.00686 | 9.43E-06 |

**Supplementary Table 11: Binomial models for testing differences in heterozygosity values of immune loci vs neutral loci in each Indian subpopulation**

| **Population** | **p-value** | **Pseudo-R-squared** |
| --- | --- | --- |
| **Indian populations** |  |  |
| **CI** | 0.0191 | 3.13E-06 |
| **SI** | 3.89e-05 | 1.29E-05 |
| **NW** | 0.0485 | 2.76E-06 |

**Supplementary Table 12: Kruskal Wallis tests for testing differences in nucleotide diversity values of neutral, receptor immune and signaling immune loci across populations**

| Comparison | Type of loci | p-value |
| --- | --- | --- |
| Across subspecies | Receptor | 0.0296* |
| Across subspecies | Signalling | 0.00001066*** |
| Across subspecies | Neutral | < 2.2e-16*** |
| Across subpopulations | Receptor | 0.0889 n.s |
| Across subpopulations | Signalling | 0.0004248*** |
| Across subpopulations | Neutral | < 2.2e-16*** |

**Supplementary Table 13: Kruskal Wallis tests for testing differences in heterozygosity values of neutral and immune loci across populations**

| Comparison | Loci | p-value |
| --- | --- | --- |
| Heterozygosity for sub species | Immune | 1.36x10^-12^ |
| Heterozygosity for sub species | Neutral | < 2.2 x10^-16^ |
| Heterozygosity for sub populations | Immune | 1.79 x 10^-8^ |
| Heterozygosity for sub populations | Neutral | < 2.2 x10^-16^ |

**Supplementary Table 14: Genes present at high nucleotide diversity and heterozygosity in different populations**

| **Gene** |  | **High Nucleotide diversity** | | | | **High heterozygosity** | | | |
| --- | --- | --- | --- | --- | --- | --- | --- | --- | --- |
| atypical chemokine receptor 3 isoform X1 |  |  |  |  |  |  | SUM |  |  |
| C-C chemokine receptor type 5 |  |  | AMU |  |  | AMU |  |  |  |
| C-C chemokine receptor type 6 |  |  |  |  |  | AMU | SUM |  |  |
| C-C chemokine receptor type 7 |  |  |  |  |  | AMU |  |  |  |
| C-C chemokine receptor-like 2 |  |  |  |  |  |  | SUM |  |  |
| C-C motif chemokine 14 |  |  |  |  |  |  |  | SOC |  |
| C-C motif chemokine 24 isoform X1 |  |  |  |  |  |  | SUM |  | NW |
| C-C motif chemokine 5 precursor |  |  |  |  |  | AMU |  |  |  |
| C-C motif chemokine 13 |  |  |  |  |  | AMU | SUM |  |  |
| C-C motif chemokine 8 |  |  |  |  |  |  |  |  |  |
| C-X-C motif chemokine 7 |  |  |  |  |  | AMU |  |  |  |
| chemokine-like receptor 1 |  |  |  |  |  |  |  |  | NW |
| C-X-C chemokine receptor type 4 |  |  |  |  |  | AMU |  |  |  |
| C-X-C chemokine receptor type 6 |  |  |  |  |  |  | SUM | SOC |  |
| C-X-C motif chemokine 14 |  |  |  |  |  | AMU |  |  |  |
| Interleukin-1 receptor accessory protein isoform X2 |  |  |  |  |  |  |  |  | NW |
| Interleukin-1 receptor type 1 isoform X2 |  |  |  |  |  |  |  |  | NW |
| Interleukin-1 receptor-like 2 isoform X2 |  |  |  |  |  |  |  |  | NW |
| Leukocyte immunoglobulin-like receptor subfamily A member 6 isoform X4 |  | SOC |  | SUM |  | AMU | SUM | SOC |  |
| C-X-C motif chemokine 11 |  |  |  |  |  |  | SUM |  |  |
| Leukocyte immunoglobulin-like receptor subfamily B member 3 isoform X1 |  | SOC |  |  |  | AMU | SUM | SOC |  |
| Leukocyte immunoglobulin-like receptor subfamily B member 3 isoform X2 |  |  |  |  |  | AMU | SUM |  |  |
| Leukocyte immunoglobulin-like receptor subfamily B member 3 isoform X3 |  |  |  |  |  | AMU |  |  |  |
| Toll-Like Receptor 2 |  |  |  |  | NW | AMU |  |  | NW |
| Toll-Like Receptor 5 |  |  |  |  |  | AMU |  |  |  |
| Toll-Like Receptor 9 |  | SOC | AMU |  |  | AMU |  |  |  |

| Tumor necrosis factor alpha-induced protein 3 isoform X2 |  |  |  |  |  |  | SUM | SOC |  |
| --- | --- | --- | --- | --- | --- | --- | --- | --- | --- |
| Tumor necrosis factor ligand superfamily member 10 |  |  |  |  |  |  |  |  | NW |
| Tumor necrosis factor ligand superfamily member 11 |  |  |  |  |  |  |  | SOC |  |
| Tumor necrosis factor receptor superfamily member 10A-like |  |  |  |  |  |  | SUM |  |  |
| Tumor necrosis factor receptor superfamily member 13C |  |  |  |  |  | AMU | SUM |  |  |
| Tumor necrosis factor receptor superfamily member 14 isoform X1 |  |  |  |  |  | AMU | SUM |  |  |
| Tumor necrosis factor receptor superfamily member 19 |  |  |  |  |  |  |  | SOC |  |
| Tumor necrosis factor receptor superfamily member 19L isoform X1 |  |  |  |  |  |  | SUM |  | NW |
| Tumor necrosis factor receptor superfamily member 21 |  | SOC |  | SUM |  |  |  |  |  |
| Tumor necrosis factor receptor superfamily member 3 |  |  |  |  |  |  |  |  | NW |
| Tumor necrosis factor receptor superfamily member 5 isoform X1 |  |  |  |  |  |  | SUM |  |  |
| Tumor necrosis factor receptor superfamily member 6 isoform X1 |  |  |  |  |  |  | SUM |  | NW |
| Tumor necrosis factor receptor superfamily member 8 isoform X4 |  |  |  |  |  | AMU |  |  |  |
| Tumor necrosis factor receptor superfamily member 9 isoform X2 |  |  |  |  | NW |  |  |  | NW |

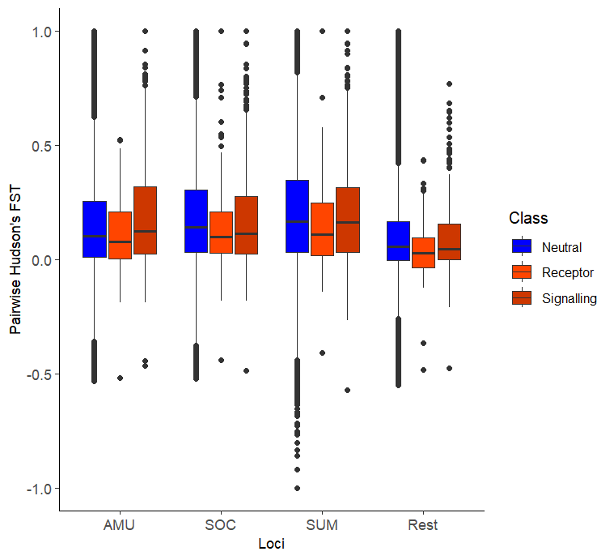

**Supplementary Figure 1: F_ST_ differences for populations of interest of tiger subspecies**

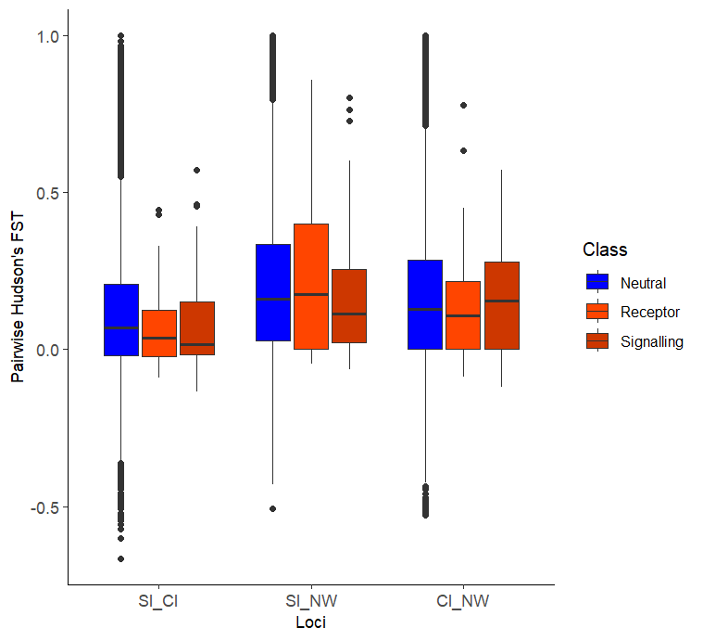

**Supplementary Figure 2: F_ST_ differences for populations of interest**

**Supplementary Table 15: Gaussian models for testing differences in F_ST_ values of immune loci versus neutral loci for each population comparison**

| **Type of loci** | **Pseudo-R Squared** | **p-value** |
| --- | --- | --- |
| **Subspecies comparison** | | |
| **AMU** |  |  |
| **Neutral** |  |  |
| **Receptor** | 4.45468E-06 | 0.00255 |
| **Signalling** |  | 5.23e-06 |
| **SUM** |  |  |
| **Neutral** |  |  |
| **Receptor** | 5.93748E-06 | 2.71e-05 |
| **Signalling** |  | 0.0748 |
| **SOC** |  |  |
| **Neutral** | 4.05474E-06 |  |
| **Receptor** |  | 2.21e-05 |
| **Signalling** |  | 0.0135 |
| Subpopulation comparison | | |
| NW vs CI |  |  |
| **Neutral** | 0 |  |
| **Receptor** |  | 0.683 |
| **Signalling** |  | 0.725 |
| NW vs SI |  |  |
| **Neutral** | 0 |  |
| **Receptor** |  | 0.2633 |
| **Signalling** |  | 0.0245 |
| SI vs CI |  |  |
| **Neutral** | 3.1294E-05 |  |
| **Receptor** |  | 0.1745 |
| **Signalling** |  | 0.0649 |

**Supplementary Table 16: Binomial Generalized Linear models for testing differences in site frequency spectra of synonymous versus neutral loci as well as nonsynonymous versus neutral SNPs**

| **Comparison** | **p-value** |
| --- | --- |
| **All immune genes** | |
| **Synonymous vs Neutral** | 1 |
| **Nonsynonymous vs Neutral** | 1 |
| **TLR** | |
| **Synonymous vs Neutral** | 1 |
| **Nonsynonymous  vs Neutral** | 1 |
| **LILR** | |
| **Synonymous vs Neutral** | 1 |
| **Nonsynonymous vs Neutral** | 1 |

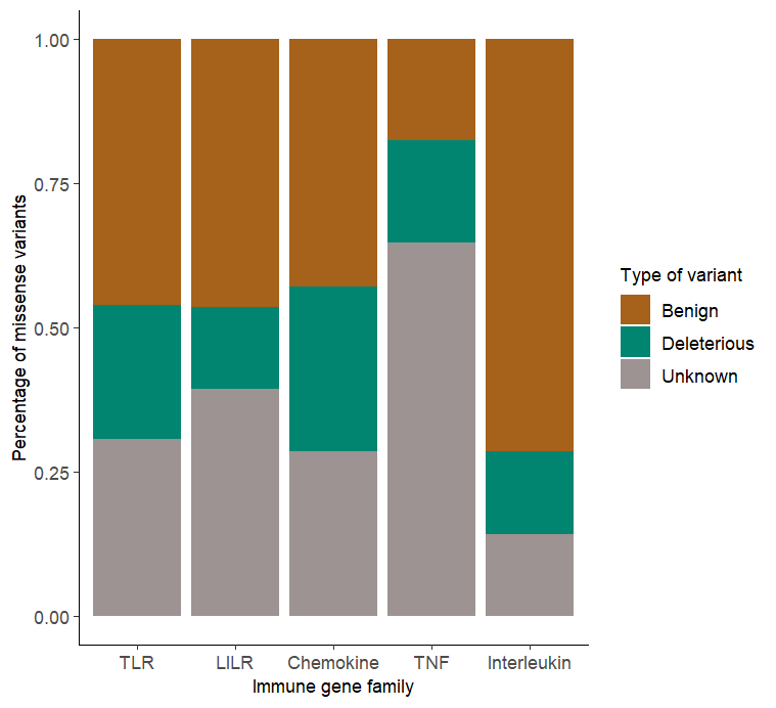

**Supplementary Figure 3: Polyphen results**
